# mRNA secondary structure stability regulates bacterial translation insulation and re-initiation

**DOI:** 10.1101/2020.02.10.941153

**Authors:** Yonatan Chemla, Michael Peeri, Mathias Luidor Heltberg, Jerry Eichler, Mogens Høgh Jensen, Tamir Tuller, Lital Alfonta

**Author notes:** Co-first authors (Yonatan Chemla, Michael Peeri).

## Abstract

In bacteria, translation re-initiation is crucial for synthesizing proteins encoded by genes that are organized into operons. The mechanisms regulating translation re-initiation remain, however, poorly understood. We now describe the ribosome termination structure (RTS), a conserved and stable mRNA secondary structure precisely localized downstream of stop codons, which serves as the main factor governing re-initiation efficiency in a synthetic *Escherichia coli* operon. We further report that in 95% of 128 analyzed bacterial genomes representing all phyla, this structure is selectively depleted when re-initiation is advantageous yet selectively enriched so as to insulate translation when re-initiation is deleterious.

## Introduction

In bacterial operons, the intergenic distance between most pairs of neighboring cistrons is shorter than 25-30 nucleotides^1,2^, a distance too small to simultaneously accommodate one ribosome terminating on the stop codon of the proximal gene and a second ribosome initiating *de novo* translation on the start codon of the distal gene^2^. Translation re-initiation alleviates this problem, as the terminating proximal-ribosome does not dissociate from the mRNA after termination and instead re-initiates translation on the neighboring distal cistron. Presently, the mechanisms regulating translation re-initiation are not well understood^2,3^. Specifically, regulators that determine whether a ribosome dissociates from the mRNA or re-initiates translation have yet to be discovered. We thus considered whether mRNA secondary structure could serve this role, given how mRNA structure can affect translation at the *de novo* initiation^4,5^ and elongation^6,7^ stages.

## Results

### mRNA structure drives distal gene expression

To test for a relation between mRNA secondary structure and translation re-initiation, a library of operons based on the pRXG plasmid^8^ was assembled (Fig. 1a). These synthetic operons comprise a proximal gene encoding red fluorescent protein (RFP) and a distal gene encoding polyhistidine-tagged green fluorescent protein (GFP), separated by a stretch of 24 random nucleotides in the inter-cistronic region, downstream of the *RFP* stop codon. The library was transformed into *Escherichia coli* MG1655 cells and sorted according to GFP levels, which spanned three orders of magnitude (Fig. 1b), into 8 bins using flow cytometry (Fig. 1c). Each bin was barcoded, sequenced, and the average Gibbs free energy (ΔG_fold_) of mRNA secondary structure in the variable sequence region was calculated (Fig. 1d). The results revealed a correlation between the observed GFP levels and the calculated mean ΔG_fold_ of the ∼3×10^3^ unique sequences in each bin. This translates to an inverse correlation, whereas in all the bins that express more than negligible levels of GFP (bins P4-P8), as compared with the negative control (fig. S1), the expression levels of the distal GFP increases as the intergenic mRNA folding stability decreases.

**Fig. 1.**
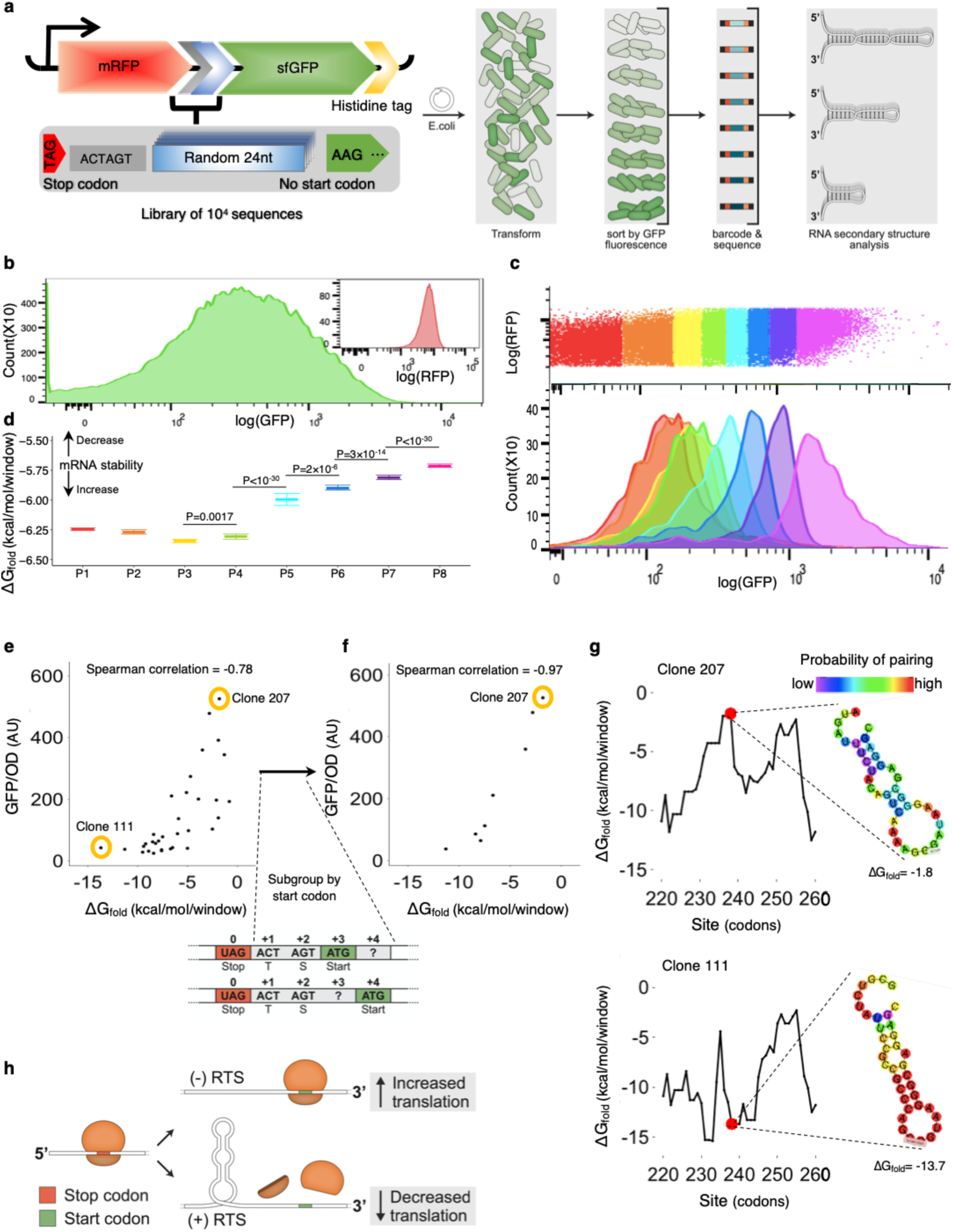
mRNA secondary structure (ΔG_fold_) controls distal operon gene expression. (**a**) Synthetic operon design and FACS sorting scheme. (**b**) GFP and RFP fluorescence of 10^5^ cells. (**c**) Sorting of 10^6^ cells into color-coded bins with constant RFP and variable GFP levels (top); GFP distribution in 3,000 cells from each bin after sorting (bottom). (**d**) The weighted mean of ΔG_fold_ with 99% confidence intervals of N=∼3×10^3^ unique sequences in each bin. Significance levels by two-sided Wilcoxon test (Table S3). Correlation between GFP expression and ΔG_fold_ of (**e**) all (n=33) isolated variants, and (**f**) a subset (n=8) presenting an AUG start codon at position +3 or +4. (**g**) mRNA secondary structure and ΔG_fold_ landscape of variable sequences of two selected clones (111, 207). (**h**) Schematic depicting the role of the RTS in distal operon gene translation.

Next, individual clones from each bin were sorted and sequenced. Thirty-three clones in which the variable inter-cistronic sequence encoded at least one of the six efficient start codons for translation initiation^9^, lacked additional in-frame stop codons and presented a unique ΔG_fold_ were isolated, and their GFP expression was quantified (Table S1). Upon assessing the relation between ΔG_fold_ of the variable sequence and GFP expression, a clear correlation was revealed (Spearman correlation ρ=-0.78, n=33, p-value<10^−7^) (Fig. 1e). Such correlation was independent of mRNA abundance (fig. S2), expression of the upstream *RFP* gene (fig. S3) or location or identity of the start codon and adjacent Shine-Dalgarno (SD) sequence in the downstream *GFP* gene to which the ribosome binds^10^ (Table S2). In a distinct subset of clones with an AUG *GFP*-start codon three or four codons downstream of the *RFP* stop codon with the same SD sequence, such correlation was strengthened (Spearman correlation ρ=-0.98, n=8, p-val=4×10^−4^) (Fig. 1f). The results thus show that distal operonic GFP gene expression is negatively affected by a stable mRNA secondary structure in the region directly downstream of the proximal stop codon (Fig. 1g), termed the ‘ribosome termination structure’ (RTS), with the likelihood of RTS presence and its strength being defined by the magnitude of ΔG_fold_ (Fig. 1h).

### The RTS is conserved across bacterial genomes

To assess the generality of the RTS, mRNA secondary structure stability (ΔG_fold_) was calculated in a region spanning 100 nucleotides on either side of each of the ∼4,200 annotated *E. coli* stop codons using a 40 nucleotide-long sliding window, allowing for calculation of a genome-wide mean ΔG_fold_ at each position (Fig. 2a). Such analysis revealed an extreme drop in ΔG_fold_ (reflecting stronger mRNA folding), with a global minimum centered 9 nucleotides downstream of stop codons (Fig. 2b, blue line), corresponding to the expected position and magnitude of an RTS. As such, RTS-like signals are apparent throughout the *E. coli* genome.

**Fig. 2.**
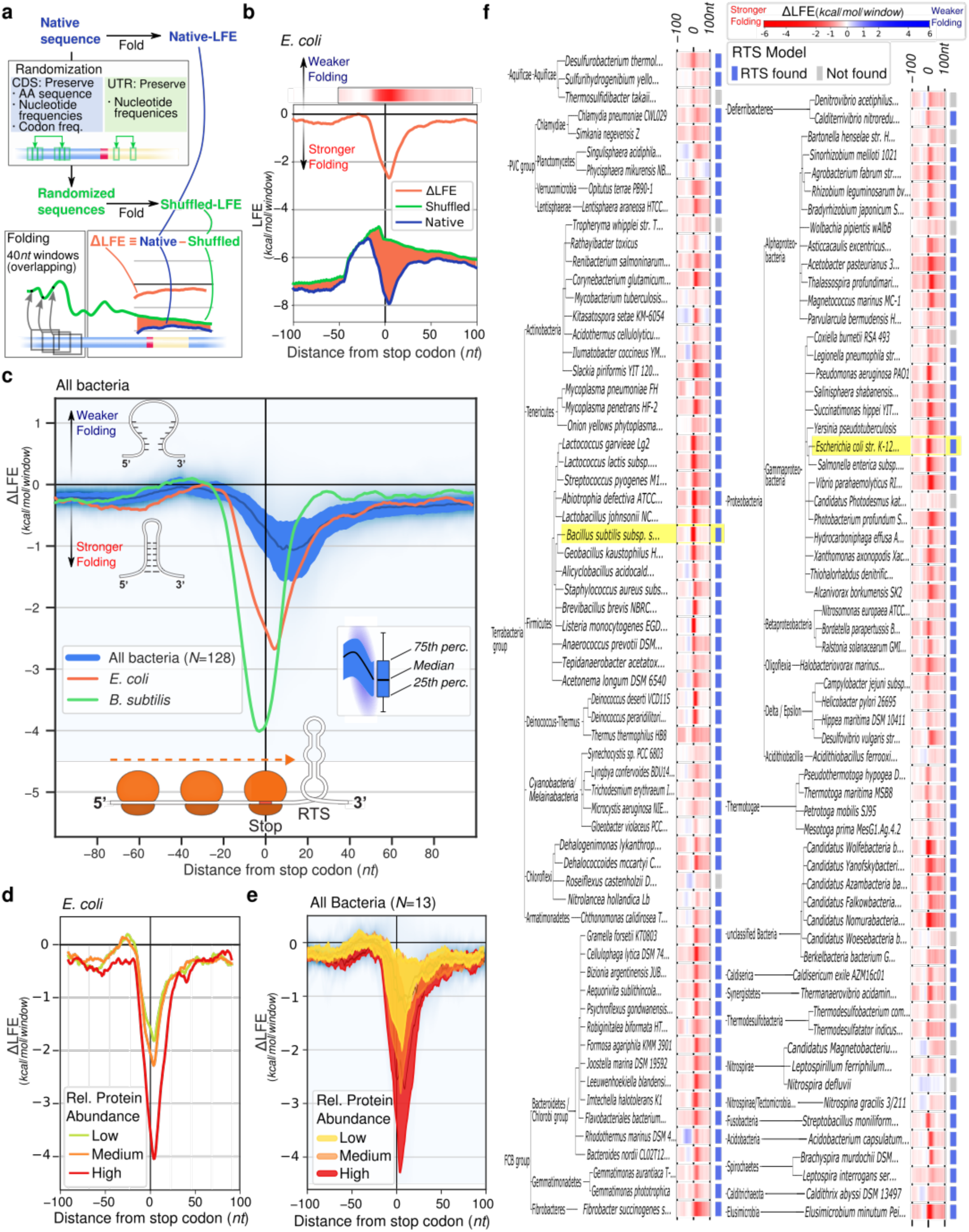
RTSs are conserved across bacterial phyla. (**a**) Pipeline for genome-wide RTS analysis. ΔLFE analysis reveals that, on average, an RTS is present and localized downstream of stop codons across all (**b**) *E. coli* and (**c**) *B. subtilis* genes and an average of 128 bacterial species examined. The RTS signal is more significant in genes encoding highly abundant products in (**d**) *E. coli*, and (**e**) all bacterial species for which protein abundance data is available. (**f**) ΔLFE heatmap depicting the 100 nucleotide-long regions around stop codons across bacteria (warm colors: stronger folding than expected; cool colors: weaker folding than expected).

To confirm that the RTS is directly under selection, the ΔG_fold_ value of each sequence (Fig. 2b, blue line), less the ΔG_fold_ value of a shuffled version in which nucleotide and codon content but not their order are preserved, was calculated (Fig. 2b, green line). Repeated for each position across all *E. coli* genes, this provided an average selection landscape of mRNA structure (Fig. 2b, orange line). If only nucleotide or codon content were under selection, then the difference in local folding energy (ΔLFE) between the native and randomized sequences should equal zero. Hence, increased deviation in the negative direction indicates direct selection for enhanced secondary structure stability (and vice versa). The results revealed extreme selection for stable structure directly downstream of stop codons (Fig. 2b, orange line) (Wilcoxon test, p-val<10^−30^), irrespective of the stop codon used (fig. S4). The same signal was seen in an average of 128 other bacterial strains representing all phyla (Fig. 2c, blue line), including the evolutionary distant Gram-positive *Bacillus subtilis* (Fig. 2c, red line). Furthermore, if RTS presence is under selection, correlation to the level of gene expression would be expected, with genes encoding more abundant proteins being subjected to stronger selection pressure. To test this hypothesis, *E. coli* genes were grouped according to protein abundance, and the ΔLFE landscape of each was determined (Fig. 2d). Clear and significant correlation protein abundance and ΔLFE was noted (Mann-Whitney test, p-val<10^−30^), demonstrating the RTS to be an adaptive trait, possibly controlling distal operon gene translation. This relation also holds true in *B. subtilis* and all 11 other bacteria for which data is available (Fig. 2e). Lastly, RTS presence was quantified genome-wide across bacteria. This revealed that RTSs, defined by an mRNA structure directly downstream of the stop codon that is significantly more stable than the surrounding sequences (see Methods), are present in 95% of the bacterial strains considered (Fig. 2f, fig. S5).

### Translation re-initiation is controlled by RTS

The precise role of the RTS was considered by examining variability in ΔLFE, distinguishing between genes followed by an RTS or not. Such analysis showed the standard deviation of ΔLFE to spike at the stop codon (Fig. 3a), yielding a bi-modal pattern of gene distribution only around the stop codon (Fig. 3b). The parameter best defining the two groups of gene distribution is the inter-cistronic distance separating neighboring genes (Fig. 3b, inset). *E. coli* gene pairs separated by shorter distances (<25 nucleotides, N=1,537) were significantly depleted of RTSs (mean ΔLFE = +0.4 kcal/mol, Wilcoxon test, p-val=5×10^−19^); for further separated adjacent genes (≥25 nucleotides, N=2,581), RTSs were significantly enriched (mean ΔLFE = −4.0 kcal/mol, Wilcoxon test, p-val<10^−30^). When the ΔLFE landscape around the stop codon between gene pairs in each group was charted (Fig. 3c), we noted RTS absence when the intergenic distance is short, or when the two consecutive cistrons overlap. Conversely, when the intergenic distance exceeds 25 nucleotides, an RTS is present (Mann-Whitney, p-val<10^−30^). This trend is conserved in 128 bacterial species analyzed (Fig. 3d)_·_ Considering that ∼25 nucleotides are the intergenic distance below which translation re-initiation is considered to be advantageous over *de novo* initiation^2^, and the above-identified correlation between RTS presence and expression of the distal operonic *GFP* gene (Fig. 1), the RTS can be linked to translation re-initiation.

**Fig. 3.**
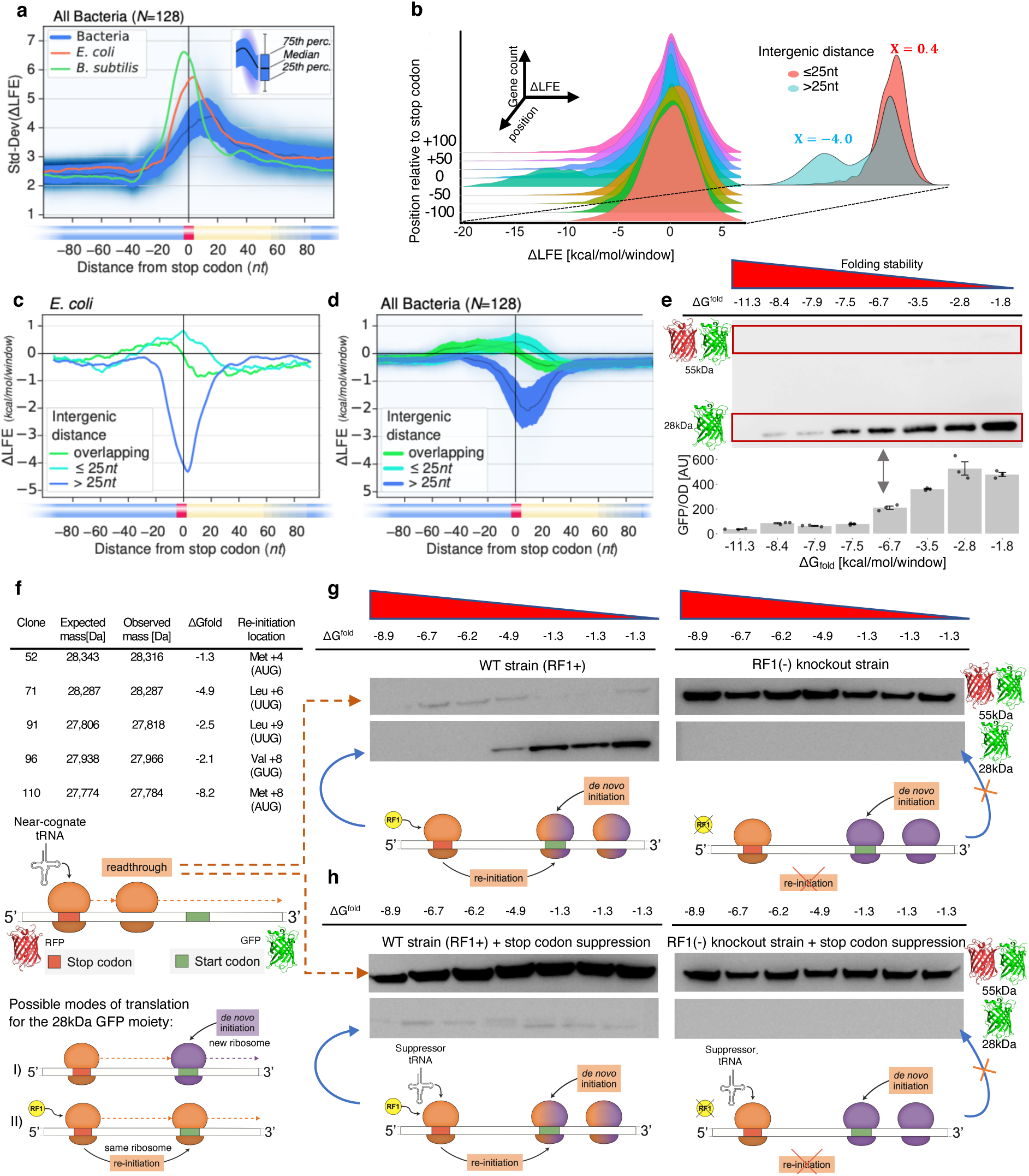
RTS is a translation re-initiation regulator. (**a**) ΔLFE standard deviation landscape around the stop codon. (**b**) *E. coli* gene density plot (Z-axis) versus ΔLFE (X-axis) and distance from a stop codon (Y-axis). Inset shows gene density at position zero. The RTS profile around the stop codon depends on the inter-cistronic distance before the downstream gene in (**c**) *E. coli* and (**d**) 128 bacterial species. (**e**) Representative anti-His-tag Western blot (top) and n≥3 fluorescence measurements with SE error bars (bottom) of eight AUG (+3/4) clones, with ΔG_fold_ indicated. (**f**) Mass spectrometry analysis of GFP from selected library clones, and the codon and location used for re-initiation. Representative and cropped Western blots of the same seven random clones from *E. coli* (**g**) without or (**h**) with stop codon reassignment, each in the presence (left) or absence (right) of RF1. Uncropped blots are available (Fig. S8)

Translation of the distal partner of any operon-based gene pair can be realized by *de novo* initiation, translation re-initiation, or stop codon read-through. Thus, discounting a link between the RTS and *de novo* initiation or stop codon read-through would further support a role for the RTS in translation re-initiation. Accordingly, experiments involving the synthetic operon described above (Fig. 1a) were performed, given how the distal GFP expression could result from any of the above-mentioned processes. The link between the RTS and stop codon read-through was tested by Western blot analysis of the above-mentioned subgroup of clones (Fig. 1f) expressing the RFP-GFP synthetic operon, normalized by OD_600_, using antibodies against the GFP C-terminal polyhistidine tag. The 55 kDa RFP+GFP product resulting from stop codon read-through was barely detectable, compared to the 28 kDa GFP product resulting from *de novo* initiation or re-initiation (Fig. 3e). The electrophoresis results of the clones, together with other randomly selected clones, were quantified by densitometry. This confirmed that the correlation between the level of the 28 kDa product and ΔG_fold_ was maintained (Spearman correlation ρ=0.80, n=58, S=6,479 p-val<10^−13^) (fig. S6). Lastly, exact product masses were verified by mass spectrometry to reveal the initiation codon and its location (Fig. 3f, fig. S7, Table S1). These findings thus discount a linkage between RTS presence and stop codon read-through.

To determine whether the RTS is linked to *de novo* initiation or translation re-initiation, the manner of *GFP* translation initiation was assessed using the release factor 1 (RF1)-deficient *E. coli* C321.Δ*prfA* EXP strain^11^ and Western blot analysis of random clones, as above. In the absence of RF1, the ribosome cannot efficiently terminate translation at the *RFP* UAG stop codon, thereby precluding translation re-initiation, which depends on such termination. Instead, GFP expression can only be driven by read-through or *de novo* initiation in the mutant strain. As Western blot analysis detected only the read-through RFP+GFP product (Fig. 3g, fig. S8), *de novo* initiation does not drive GFP translation. Still, the apparent lack of *de novo* GFP translation initiation in the deletion strain could result from physical interference of the initiation site by *RFP*-translating ribosomes and increased read-through. To discount this possibility, the *RFP* UAG stop codon in *E. coli* MG1655 was suppressed (see Methods) so as to mimic conditions of ribosomal occupancy that may occur in RF1-deficient cells. As also under these conditions, isolated GFP was produced only in the *E. coli* MG1655 strain but not in RF1-depleted cells (Fig. 3h); *de novo* initiation could be excluded, leaving translation re-initiation as the only conceivable process by which RTS regulates expression of the operonic distal *GFP* gene.

### RTS is dependent on gene’s operon-position

Finally, to determine whether the translation re-initiation regulator role assigned to RTS can be generalized, “transcriptional unit” data^12^ cataloging the arrangement of *E. coli* genes into operons was assessed (Fig. 4a). After all operon terminal genes, where re-initiation is deleterious, the presence of an RTS following the stop codon is favored. In contrast, RTSs are depleted following the stop codon of all other operonic genes, thus encouraging re-initiation (Mann-Whitney, p-val<10^−30^). These results were strengthened by observing that RTS presence after terminal genes is independent of the presence or absence of start codons in the 50 nucleotide-long stretch downstream of the stop codon, while significant dependence was seen for other operon genes (fig. S9). The same held true in *B. subtilis* and four other bacterial species for which experimental operon arrangement data exists (Fig. 4a). Finally, gene annotations for 128 bacterial species were analyzed for RTS presence as a function of neighboring gene strand directionality. Accordingly, pairs of neighboring genes on the same strand, where re-initiation on the mRNA is possible, were compared to pairs on opposite strands, where such re-initiation would be useless as the opposite annotation could not be translated on the same mRNA (Fig. 4b). As expected, RTS presence was significantly higher within gene pairs found on opposite strands. These results, together with the demonstrated lack of RTS linkage to transcription termination (figs. S2&S10), are all consistent with the RTS being a general regulator of translation re-initiation in bacteria.

**Fig. 4.**
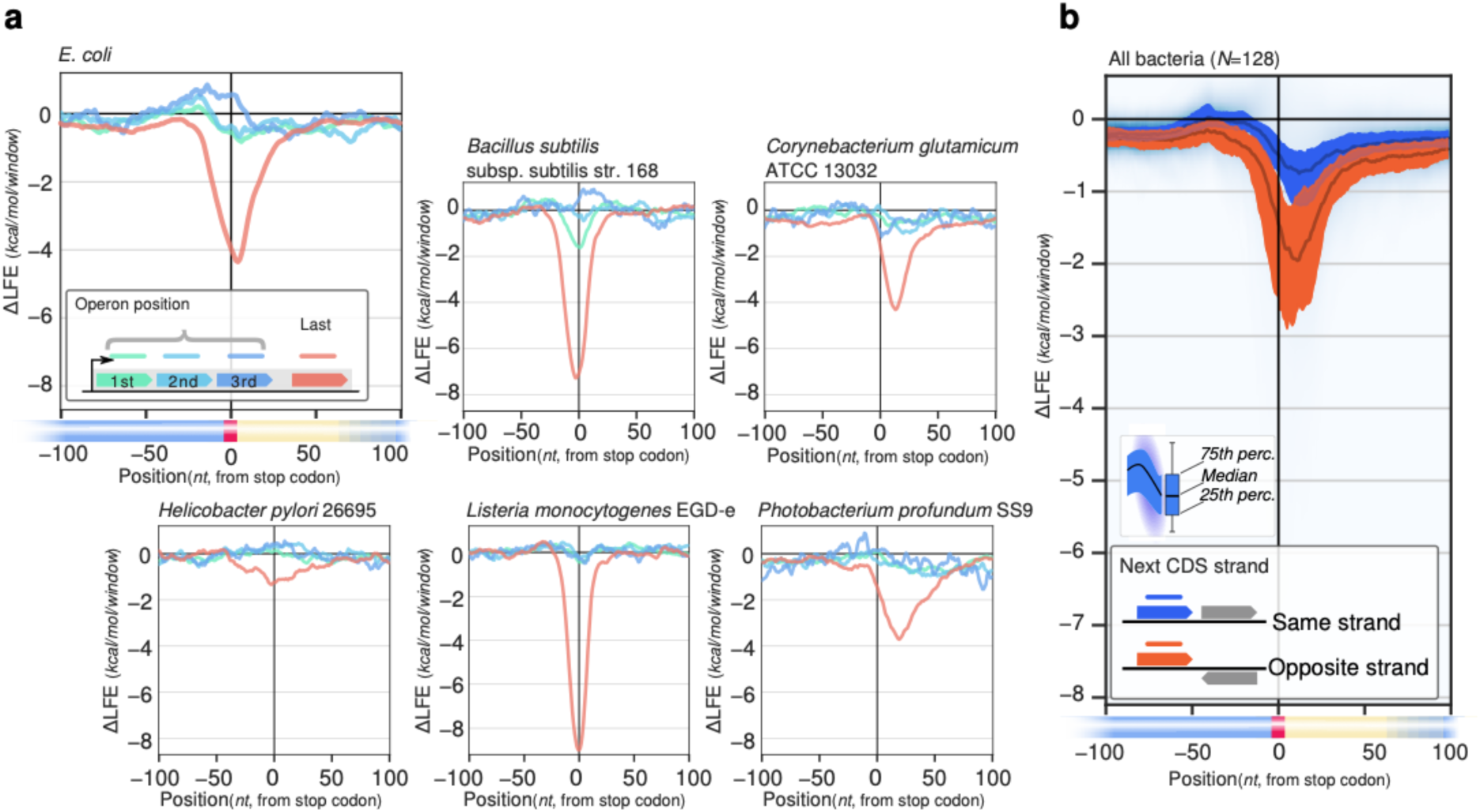
In all bacteria phyla, RTSs are enriched where re-initiation is deleterious and depleted where re-initiation is advantageous. (**a**) RTS presence depends on operonic position in *E. coli* and in all operon-mapped bacterial species. (**b**) RTS presence depends on the directionality of the downstream cistron in 128 bacterial species.

## Discussion

Translation re-initiation affords bacteria the ability to translate operon-sequestered genes without significant interference between terminating and initiating ribosomes. However, translation re-initiation also carries risk. Uncontrolled, re-initiated translation could evoke high fitness costs due to ribosomes devoting more time scanning for re-initiation or because of unintended translation re-initiation events. Indeed, as the ribosome can re-initiate in all possible frames and recognizes several start codons and alternative SD sequences (Tables S1&S2), unintended translation re-initiation is of real concern (fig. S11). As such, regulation of translation re-initiation is needed.

Here, we identified a stable mRNA secondary structure downstream of the stop codon (termed the RTS) that controls translation re-initiation. We revealed that robust signals corresponding to the presence of an RTS are found across the *E. coli* genome, in agreement with recently published transcriptome-wide mRNA stability data^13,14^. We also showed the RTS to be conserved across bacterial phyla, with an RTS signal peaking at a position that correlates with the edge of the mRNA stretch that is shielded by a terminating ribosome, alluding to a possible RTS-ribosome interaction. The functional analyses and experiments performed here all support the RTS acting as a translational insulator, inhibiting translation re-initiation. We cannot exclude that the RTS also serves as an inhibitor to *de novo* initiation. However, we did not observe such initiations, and model predictions did not correlate with our results (Table S2). In addition, the expression of upstream RFP was independent of the strength of expression of the downstream GFP (fig. S3). The latter is unexpected when GFP translation is *de novo*-initiated, as the distance between the RFP stop codon and the GFP start codon is too small (6-24 nucleotides) to allow both genes to bear terminating and initiating ribosomes, simultaneously. Accordingly, these ribosomes must compete and be inter-dependent. The expression of both genes appears, however, to be independent, thus strengthening the conclusion of the RTS insulating re-initiation.

Currently, two competing models explain re-initiation, namely the classic 30S-binding model, where ribosomes dissociate from polycistronic mRNA upon gene translation termination only to immediately re-bind, as in *de novo* initiation and translate the downstream cistron ^10^. In this mode, one would expect to detect translation of a distal cistron by both re-initiating and *de novo* initiating ribosomes, which would compete for the ribosome binding domain. The second model is the recently demonstrated 70S-scanning model, where the ribosome does not dissociate but instead scans downstream mRNA for a re-initiation site^2^. Our results provide support for the latter model as *de novo* initiation was not observed. Moreover, the observed existence of an RTS in terminal genes is more parsimonious when scanning-based re-initiation occurs. Although the molecular mechanism by which the RTS controls ribosomal re-initiation remains unknown, we can conjecture, given earlier reports, that it acts as an energy barrier for the scanning ribosome, which unlike the actively elongating ribosome, does not have an energy source^2,15,16^. In summary, the discovery of the ribosome termination structure, a translation re-initiation insulator, raises new questions on the function and evolution of operons and could lead to exploitation of the remarkably conserved RTS trait for better control over genetic design.

## Methods

### Experimental methods

#### Strains and plasmids

The bacterial strains used in this study were *Escherichia coli* K-12 MG1655 (Yale stock CGSC#: 6300) and *E. coli* C321.Δ*prf*A EXP^11^ (Addgene #48998). For stop codon suppression by genetic code expansion, experimental strains were transformed with a pEVOL plasmid harboring the *Methanosarcina mazei* (*Mm*) orthogonal pair of *Mm*-PylRS/*Mm*-tRNA_CUA_^PrK^ (Pyl-OTS)^17,18^. The synthetic operon plasmid was adapted from the pRXG dual reporter plasmid^8^, and the random sequence was inserted using random primer amplification followed by Gibson assembly. The expression of the synthetic operon was controlled by the Lac operator as to not affect bacterial fitness by the variability of the random sequence, which is only expressed when IPTG is supplemented (1mM) to the growth media. To control for known stop codon context effects^19^, the first six nucleotides in this variable region (ACUAGU) were fixed. After assembly, the library was transformed into *E. coli* DH5α, where library complexity was measured to be ∼10^4^ by counting colony-forming units. The plasmid library was then purified using a Miniprep kit [Promega] and transformed into the *E. coli* MG1655 and C321 strains mentioned above. All *E. coli* MG1655 clones were subjected to fluorescence-activated cell sorting (FACS) [FACSAria, BD Biosciences]. In addition, individual clones were isolated using agar plating, and their plasmids purified and sequenced. Each variable sequence that did not present an additional stop codon in the variable region was named pRXNG and given a running number name (i.e., pRXNG 60 is clone #60) and its RFP and GFP expression levels were measured.

#### Fluorescence-activated cell sorting

Bacterial cells were grown overnight induced with 1 mM IPTG, washed with PBS, and sorted by using FACS [FACSAria III, BD Biosciences]. The entire cell population was sorted into 8 bins based on constant mRFP1 fluorescence and varying Superfolder GFP (sfGFP) fluorescence, thereby normalizing sfGFP levels to those of mRFP1. Each bin accounted for ∼12.5% of the entire population, using an 85-micron nozzle at minimal flow. The 8 sorted bins were re-run to map sorting accuracy, which was found to be high (∼90% of cells were distributed within 3 bins around any selected bin). Controls consisted of bacterial cells that did not harbor the synthetic operon plasmid. The analysis was done, and figures created using the FlowJo software. The gating strategy was as follows: The preliminary FSC-A/SSC-A gates were: 630-17,000 and 60-3,000 respectively, The SSC-W/SSC-H gates were 0-110,000 and 450-45,000, respectively, and the FSC-W/FSC-H gates were 12,000-62,000 and 200-4,000, respectively. Cells that expressed RFP which served as the positive and normalizing control with levels between 3,500-15,000, were further gated. Next, the resulting population (49.7% of the total population) was gated into 8 ∼equal groups divided and defined by GFP expression; each group was intended to represent ∼12.5% of the parent population.

#### Library construction, next-generation sequencing, and data analysis

Isolated bacteria from each bin were transferred to LB media and grown for 8h at 37°C. Cells were harvested and subjected to plasmid extraction using a Miniprep kit [Promega]. Library construction for Illumina MiSeq next-generation sequencing was done under the Illumina metagenomic protocol^20^. In each bin, a 118bp synthetic operon amplicon, which includes the variable region, was PCR-amplified. In two rounds of amplification, the Illumina primer sequence, unique hepta-nucleotide indexes, and adaptors were added to each amplicon library. The libraries were then sequenced using the Illumina MiSeq V2 reagent (300 cycles) kit. The resulting sequencing data was processed and parsed with the DADA2 package for R^21^. All identical sequence reads in each bin were aggregated, and the 10,000 most abundant sequences of each bin were obtained. In the eight bins, the minimal sequence depth was 2-10 reads. From the 10,000 unique sequences of each bin, all sequences which contained an additional stop codon in the variable region were removed, and the remaining sequences were filtered to include only sequences with one of the three efficient start codons (ATG, GTG, TTG) ^9^ in any in-frame position of the variable region. This process resulted in N=2,580-2,694 unique sequences in each bin. The mean ΔG_fold_ and the 99% confidence interval were calculated for each bin (see computational method for calculation) and the statistical significance comparing each pair of consecutive bins was done using a two-tail Wilcoxon rank test.

#### RFP and GFP expression from the dual reporter with the random library

Measurements from triplicate bacterial growth cultures in a 96-well plate [Thermo Scientific] covered with Breathe-Easy seals [Diversified Biotech] were recorded overnight using a 37°C incubated plate reader [Tecan]. RFP (excitation: 584 nm; emission: 607 nm) and sfGFP (excitation: 488 nm; emission: 507 nm) expression levels and OD_600_ were measured every 15 minutes. The values presented the plateau value of each clone, which was measured in at least 3 experimental repeats (n≥3).

#### Western blots

Bacterial cultures were normalized to the same OD_600_, after which 10 μL aliquots were mixed with 10 μL MOPS buffer and 5 μL SDS buffer and incubated for 10 min at 70°C. Samples were loaded onto a 4-20% SDS gel [Genscript] and transferred to a PVDF membrane [Bio-Rad] using an E-blot protein transfer apparatus [Genscript]. After transfer, anti-His tag antibodies [his-probe (H-3) antibodies, Santa Cruz Biotechnology, sc-8036, Lot #B2317] were used to probe the transferred proteins. Antibody binding was visualized using an ImageQuant LAS 4000 imager [Fujifilm]. Densitometry analysis was done using the gel tool in the ImageJ V1.52a software.

#### Stop codon suppression by genetic code expansion

Genetic code expansion by stop codon suppression was introduced to suppress the UAG stop codon in *E. coli* MG1655, where the unnatural amino acid N-propargyl-l-lysine (1 mM final concentration in culture) was incorporated in response to the UAG stop codon at the end of the RFP gene using the *Mm* pyrrolysine tRNACUApyl and pyrrolysyl-tRNA synthetase orthogonal pair^22^, expressed from the pEVOL plasmid ^17,18^. Induction of PylRS was performed by adding 0.5% L-arabinose [Sigma-Aldrich] to the growth medium.

#### Quantitative PCR

*E. coli* MG1655 cells were grown to logarithmic phase, harvested, and treated with a GeneJET RNA purification kit [Thermo Scientific] for total mRNA extraction. The RNA was immediately reverse-transcribed into cDNA by the iScript cDNA Synthesis kit [Biorad]. Real-time PCR was performed using a KAPA SYBR FAST qPCR kit [KapaBiosystems] with the recommended relative calibration curve protocol in a StepOnePlus Real-Time PCR System [Thermo Scientific].

#### Protein purification and mass spectrometry analysis

Proteins were fused to a 6xHis tag and purified by nickel resin affinity chromatography. Purified protein samples were analyzed by LC-MS [Finnigan Surveyor/LCQ Fleet, Thermo Scientific].

### Computational methods

#### Calculation of ΔG_fold_ for synthetic operon clones

All calculations were made using the Vienna package^23^ (default settings), with the extracted mRNA sequence window upon which the ΔG_fold_ calculation was made for each clone obeying the two following constraints: First, the start of the window was +9 nucleotides from the first nucleotide of the UAG stop codon. This was done to simulate mRNA secondary structure, which exists outside the ribosomal entry tunnel. Second, the window size used was experimentally determined, with a threshold requirement, namely correlation between ΔG_fold_ and GFP expression, should be robust using window sizes ranging from 30 to 50 nucleotides (length of the random region of interest = 24 nucleotides). The optimal correlation was found with a window size of 37 nucleotides. As such, this window size was used for the results presented.

#### Species selection

Species were chosen for taxonomic diversity and overlapped with public datasets (*N*=183), with emphasis on bacteria (N=128) and archaea (N=49). Genomic sequences and annotations were obtained from the Ensembl database^24^.

#### ΔLFE (folding bias) calculations

To estimate the tendency of short-range interactions within the mRNA strand to form stable secondary structures, i.e., Local Fold Energy (LFE), sequences were broken into 40 nucleotides-long windows, and the minimum folding energy was calculated using RNAfold from the Vienna package^23^ (using default settings). To identify regions where strong or weak secondary structure may be functional, rather than a side effect of selection acting on amino acid sequence, or nucleotide or codon composition (see Randomization, below), the influence of these factors was controlled by comparing LFE of the native sequence to a set of randomized sequences maintaining these factors. The difference between the LFE of the native and randomized sequences is denoted as ΔLFE or local folding bias. If only the amino acid sequence, nucleotide composition, and codon composition are under selection at a given position, one expects ΔLFE to be close to 0. Any statistically significant deviation from this value indicates that additional factors maintained under selection are needed to explain the measured native LFE value.

Since this study focused on mRNA, only those regions surrounding protein-coding genes are included; genes shorter than 40 nucleotides were excluded. Genes with a length that is not a multiple of 3, those containing an internal stop codon or where the last codon is not a stop codon were also excluded. To identify features related to translation termination, ΔLFE for all included genes from a given species was averaged at each position relative to the stop codon.

#### Randomization

The randomized sequences were sampled from the distribution representing the null hypothesis, namely that only the amino acid sequence, and nucleotide and codon composition (see below) are under selection at a given position in the coding sequence, and only the nucleotide composition is under selection in a given UTR. To produce random sequences maintaining these properties, synonymous codons within each coding sequence were randomly permutated, and the nucleotides of each UTR were randomly permutated. Regions overlapping multiple coding sequences were maintained without permutations. Codons containing one or more ambiguous nucleotides (‘N’ bases) were likewise maintained without permutations. Synonymous codons were identified according to the gene translation table for each species.

#### RTS model

To estimate the number of species likely to contain the RTS in many genes, ΔLFE for all genes at each position was compared to the mean folding bias at the UTR for the same species, measured as the average at positions 50 to 100 nucleotides (relative to the end of the stop codon). An RTS was deemed present if in at least 3 consecutive windows in the range −5 to 20 nucleotides (relative to the end of the stop codon), the following 2 conditions were met: 1. The mean ΔLFE at this position is stronger (more negative) than the mean UTR level; 2. ΔLFE for genes at that position is significant, relative to the mean UTR level, as determined by a Wilcoxon signed-rank test.

#### Plotting

Distributions of multiple genes or averages for multiple species are presented using the statistics commonly used for boxplots, as follows. The shaded region spans the 25th and 75th percentiles, with the median plotted as a darker line. Elements outside this region are presented by their density (blue shading in the background). Densities are shown as kernel density estimates (KDEs), computed separately at each position, using a Gaussian kernel with a bandwidth of 0.5. Plots were created using Scikit Learn^25^ and Matplotlib^26^. Taxonomic trees are based on NCBI taxonomy^27^ and were plotted using the ete toolkit^28^.

#### Statistical analysis

All statistical analysis was performed under the guidelines of the tests described in-text. The minimal p-value noted in the text was selected to be 10^−30^, in all cases where the precise p-value calculated was smaller (i.e., more significant), the test-statistic score is given. To test whether ΔLFE values for a one sample-group of genes are statistically different compared to a reference value (e.g., for the RTS model), the Wilcoxon signed-ranks test was used on the ΔLFE (randomized ΔG-native ΔG) values for all genes (20 randomization repetitions for each gene). To test whether ΔLFE values for two sample-groups of genes are statistically different from each other, the Mann-Whitney U test was used on the ΔLFE (randomized ΔG-native ΔG) values for all genes (with 20 randomization repetitions for each gene), as such, the test N was 20 times the number of data points of the original sample. The reported p-value and test statistic are reported for the position of the most extreme test-statistic, but the surrounding regions showed consistent and significant results. Detailed statistical parameters are available in Table S3.

#### Additional data sources

Experimentally-determined operonic positions were obtained from ODB4^29^. Protein-abundance data was obtained from PaxDb^30^. Experimentally determined 3’-UTR lengths were obtained from regulondb^31^. Termination type data for *E. coli* genes were obtained from WebGesTer^32^.

## Acknowledgments

We gratefully acknowledge Itay Algov, Dr. Anna Bakharat, Yariv Greenshpan, and Dvir Schirman for their invaluable advice and assistance.

## Author contributions

Y.C conceived, performed all experiments and wrote the manuscript, Y.C and M.P conceived and executed the computational analyses, M.H, M.H.J contributed to computational analyses and interpretations of data, J.E. contributed to the writing of the manuscript and interpretation of data, T.T conceived and supervised the computational analyses, L.A conceived and supervised experiments and wrote the manuscript.

## Data and materials availability

All data generated or analyzed during this study are included in this published article (and its supplementary information files), except for the next-generation sequencing data set and the tables for all, except *E. coli*, 127 bacteria analyzed, which will be made available on an online database by the time of the publication.

## Code availability

All custom codes used to generate the results of this paper will be available upon demand from Tamir Tuller.

## References

1. Huber, M. et al. Translational coupling via termination-reinitiation in archaea and bacteria. Nat. Commun. 10, (2019).

2. Yamamoto, H. et al. 70S-scanning initiation is a novel and frequent initiation mode of ribosomal translation in bacteria. Proc. Natl. Acad. Sci. 113, E1180–E1189 (2016).

3. Gunišová, S., Hronová, V., Mohammad, M. P., Hinnebusch, A. G. & Valášek, L. S. Please do not recycle! Translation reinitiation in microbes and higher eukaryotes. FEMS Microbiol. Rev. 42, 165–192 (2018).

4. Kudla, G., Murray, A. W., Tollervey, D. & Plotkin, J. B. Coding-Sequence Determinants of Gene Expression in Escherichia coli. Science. 324, 255–259 (2009).

5. Cambray, G., Guimaraes, J. C. & Arkin, A. P. Evaluation of 244, 000 synthetic sequences reveals design principles to optimize translation in Escherichia coli. Nat. Biotechnol. 36, (2018).

6. Tuller, T. et al. Composite effects of gene determinants on the translation speed and density of ribosomes. Genome Biol. 12, R110 (2011).

7. Gorochowski, T. E., Ignatova, Z., Bovenberg, R. A. L. & Roubos, J. A. Trade-offs between tRNA abundance and mRNA secondary structure support smoothing of translation elongation rate. Nucleic Acids Res. 43, 3022–3032 (2015).

8. Monk, J. W. et al. Rapid and inexpensive evaluation of nonstandard amino acid incorporation in Escherichia coli. ACS Synth. Biol. 6, 45–54 (2017).

9. Hecht, A. et al. Measurements of translation initiation from all 64 codons in E. coli. Nucleic Acids Res. 45, 3615–3626 (2017).

10. Kozak, M. Initiation of translation in prokaryotes and eukaryotes. Gene 234, 187–208 (1999).

11. Lajoie, M. J. et al. Genomically recoded organisms expand biological functions. Science 342, 357–60 (2013).

12. Gama-Castro, S. et al. RegulonDB version 9.0: High-level integration of gene regulation, coexpression, motif clustering and beyond. Nucleic Acids Res. 44, D133–D143 (2016).

13. Del Campo, C., Bartholomäus, A., Fedyunin, I. & Ignatova, Z. Secondary Structure across the Bacterial Transcriptome Reveals Versatile Roles in mRNA Regulation and Function. PLoS Genet. 11, 1–23 (2015).

14. Burkhardt, D. H. et al. Operon mRNAs are organized into ORF-centric structures that predict translation efficiency. Elife 6, 474–486 (2017).

15. Adhin, M. R. & D, J. Van. Scanning Model for Translational Reinitiation in Eubacteria. J. Mol. Biol. 213, 811–818 (1990).

16. Osterman, I. A., Evfratov, S. A., Sergiev, P. V & Dontsova, O. A. Comparison of mRNA features affecting translation initiation and reinitiation. Nucleic Acids Res. 41, 474–486 (2012).

17. Young, T. S., Ahmad, I., Yin, J. A. & Schultz, P. G. An enhanced system for unnatural amino acid mutagenesis in E. coli. J. Mol. Biol. 395, 361–374 (2010).

18. Chemla, Y., Ozer, E., Schlesinger, O., Noireaux, V. & Alfonta, L. Genetically expanded cell-free protein synthesis using endogenous pyrrolysyl orthogonal translation system. Biotechnol. Bioeng. 112, 1663–1672 (2015).

19. Chemla, Y., Ozer, E., Algov, I. & Alfonta, L. Context effects of genetic code expansion by stop codon suppression. Curr. Opin. Chem. Biol. 46, 146–155 (2018).

20. Amplicon, P. C. R., Clean-Up, P. C. R. & Index, P. C. R. 16s metagenomic sequencing library preparation.

21. Callahan, B. J. et al. DADA2: high-resolution sample inference from Illumina amplicon data. Nat. Methods 13, 581 (2016).

22. Srinivasan, G., James, C. M. & Krzycki, J. a. Pyrrolysine encoded by UAG in Archaea: charging of a UAG-decoding specialized tRNA. Science 296, 1459–62 (2002).

23. Lorenz, R. et al. {ViennaRNA} Package 2.0. Algorithms Mol. Biol. 6, 26 (2011).

24. Cunningham, F. et al. Ensembl 2019. Nucleic Acids Res. 47, D745–D751 (2019).

25. Pedregosa, F. et al. Scikit-learn: Machine learning in Python. J. Mach. Learn. Res. 12, 2825–2830 (2011).

26. Hunter, J. D. Matplotlib: A 2D graphics environment. Comput. Sci. Eng. 9, 90 (2007).

27. Agarwala, R. et al. Database resources of the National Center for Biotechnology Information. Nucleic Acids Res. 46, D8–D13 (2018).

28. Huerta-Cepas, J., Serra, F. & Bork, P. ETE 3: Reconstruction, Analysis, and Visualization of Phylogenomic Data. Mol. Biol. Evol. 33, 1635–1638 (2016).

29. Okuda, S. & Yoshizawa, A. C. ODB: A database for operon organizations, 2011 update. Nucleic Acids Res. 39, 552–555 (2011).

30. Wang, M. et al. PaxDb, a database of protein abundance averages across all three domains of life. Mol. Cell. Proteomics 11, 492–500 (2012).

31. Santos-Zavaleta, A. et al. RegulonDB v 10.5: Tackling challenges to unify classic and high throughput knowledge of gene regulation in E. coli K-12. Nucleic Acids Res. 47, D212–D220 (2019).

32. Mitra, A., Kesarwani, A. K., Pal, D. & Nagaraja, V. WebGeSTer DB-A transcription terminator database. Nucleic Acids Res. 39, 129–135 (2011).

