## Supplementary Figures and Tables for "mRNA secondary structure stability regulates bacterial translation insulation and re-initiation"

[4]Department of Biomedical Engineering, The Iby and Aladar Fleischman Faculty of Engineering and The Sagol School of Neuroscience, Tel Aviv University, Tel Aviv 6997801, Israel

[†] Co-first authors

### Figure S1 – Flow cytometry gating and negative control

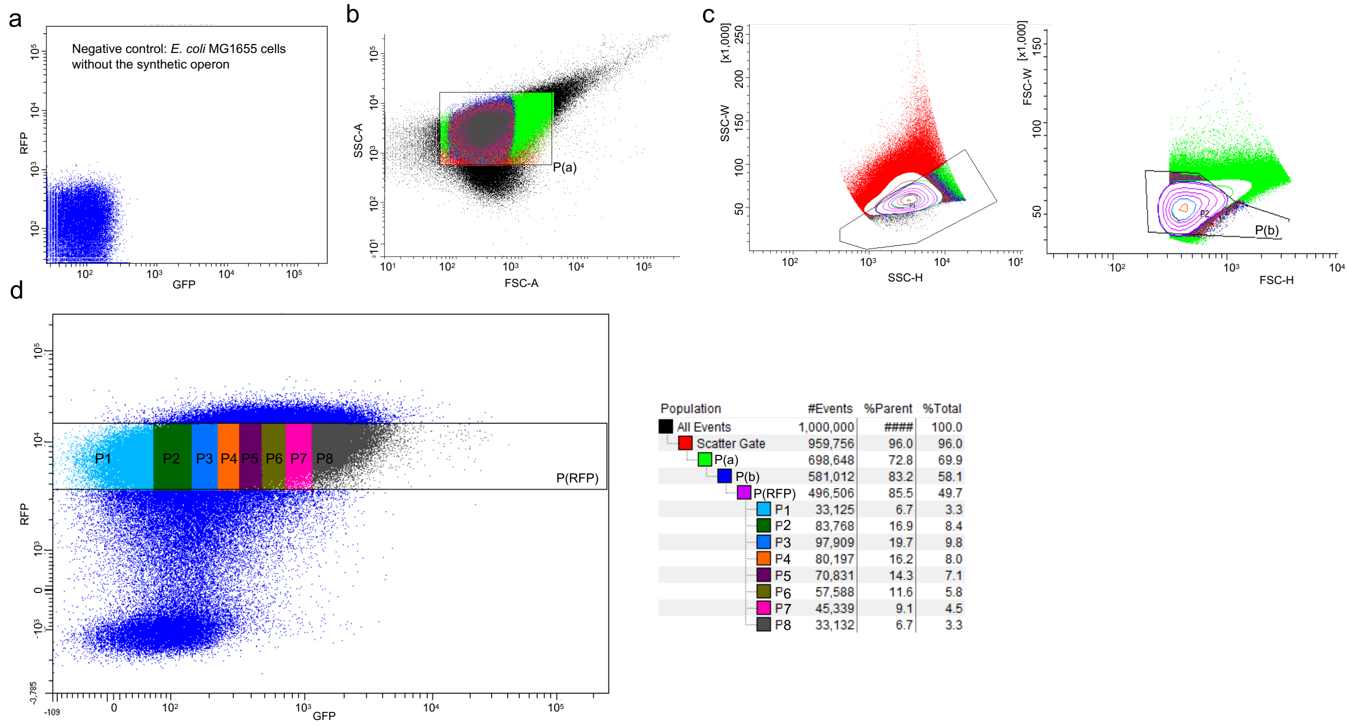


| Population | #Events | %Parent | %Total |
|---|---|---|---|
| All Events | 1,000,000 | #### | 100.0 |
| Scatter Gate | 959,756 | 96.0 | 96.0 |
| P(a) | 698,648 | 72.8 | 69.9 |
| P(b) | 581,012 | 83.2 | 58.1 |
| P(RFP) | 496,506 | 85.5 | 49.7 |
| P1 | 33,125 | 6.7 | 3.3 |
| P2 | 83,768 | 16.9 | 8.4 |
| P3 | 97,909 | 19.7 | 9.8 |
| P4 | 80,197 | 16.2 | 8.0 |
| P5 | 70,831 | 14.3 | 7.1 |
| P6 | 57,588 | 11.6 | 5.8 |
| P7 | 45,339 | 9.1 | 4.5 |
| P8 | 33,132 | 6.7 | 3.3 |

**Fig. S1.** Flow Cytometry gating and negative control. **a**) The negative control, which consists of WT *E. coli* MG1655 **b**) First size gating **c**) Second size gating **d**) uncropped sorting with gate and population statistics.

**Table S1.** Characterization of individual clones sequenced from the random library
All sequences are available as tables in the data section

| Clone | $\Delta G_{fold}$ | Average fluorescence RFP/OD [AU] ± SE | Average fluorescence GFP/OD [AU] ± SE | Average RFP/GFP | Best re-start codon | Start codon position* | Codon rank [1] | MS Verification |
|---|---|---|---|---|---|---|---|---|
| **29** | -8.9 | 697±254 | 32±4 | 18.5 | I (AUU) | +8 | 6th | |
| **33** | -9.4 | 602±174 | 48±5 | 11.7 | V (GUG) | +8 | 2nd | |
| **52** | -1.3 | 1293±269 | 344±12 | 3.8 | M (AUG) | +5 | 1st | |
| **56** | -8.5 | 624±248 | 62±4 | 9.1 | V (GUG) | +6 | 2nd | Yes |
| **57** | -6.5 | 923±416 | 43±7 | 19.1 | L (UUG) | +8 | 3rd | |
| **62** | -6.7 | 474±94 | 39±4 | 12.5 | L (CUG) | +9 | 4th | |
| **71** | -4.9 | 853±153 | 98±4 | 8.8 | L (UUG) | +6 | 3rd | |
| **91** | -2.5 | 1155±732 | 103±4 | 13.4 | L (UUG) | +9 | 3rd | Yes |
| **96** | -2.1 | 1161±452 | 197±7 | 5.8 | V (GUG) | +8 | 2nd | Yes |
| **101** | -6.0 | 496±64 | 74±24 | 6.7 | L (UUG) | +6 | 3rd | Yes |
| **110** | -8.2 | 759±228 | 57±4 | 14.8 | M (AUG) | +8 | 1st | |
| **111** | -13.7 | 486±62 | 43±3 | 10.8 | I (AUU) | +5 | 6th | Yes |
| **202** | -11.3 | 362±126 | 38±2 | 9.0 | V (GUG) | +4 | 2nd | |
| **203** | -7.6 | 320±82 | 33±6 | 9.4 | L (UUG) | +7 | 3rd | |
| **207** | -1.8 | 1236±541 | 526±53.5 | 2.1 | M (AUG) | +3 | 1st | |
| **208** | -1.9 | 1163±664 | 140±19 | 6.5 | M (AUG) | +7 | 1st | |
| **209** | -4.7 | 276±42 | 274±8 | 1.1 | M (AUG) | +7 | 1st | |
| **212** | -5.9 | 287±83 | 137±5 | 2.0 | M (AUG) | +3 | 1st | |
| **214** | -2.8 | 313±75 | 478±17 | 0.7 | M (AUG) | +3 | 1st | |
| **216** | -0.8 | 360±78 | 193±17 | 1.9 | M (AUG) | +5 | 1st | |
| **220** | -3.9 | 354±112 | 201±16 | 1.7 | M (AUG) | +6 | 1st | |
| **222** | -6.6 | 333±104 | 211±13 | 1.7 | M (AUG) | +3 | 1st | |
| **225** | -7.5 | 319±23 | 78±4 | 4.0 | L (UUG) | +3 | 3rd | |
| **226** | -9.2 | 367±24 | 56±6 | 6.8 | L (UUG) | +5 | 3rd | |
| | | | | | M (AUG) | +9 | 1st | |
| **230** | -5.0 | 320±29 | 41±6 | 8.2 | M (AUG) | +8 | 1st | |
| **232** | -7.6 | 378±34 | 36±5 | 10.3 | M (AUG) | +6 | 1st | |
| **233** | -8.4 | 398±37 | 25±5 | 18 | V (GUG) | +8 | 2nd | |
| **235** | -9.5 | 282±13 | 29±5 | 10 | L (CUG) | +5 | 4th | |
| **236** | -7.9 | 402±58 | 65±4 | 6.5 | L (UUG) | +3 | 3rd | |
| **238** | -8.4 | 367±41 | 86±5 | 4.4 | V (GUG) | +4 | 2nd | |
| **244** | -3.5 | 362±54 | 360±5 | 1 | M (AUG) | +4 | 1st | |
| **245** | -5.1 | 411±48 | 222±11 | 1.9 | M (AUG) | +7 | 1st | |
| **249** | -1.9 | 406±50 | 391±18 | 1.1 | M (AUG) | +7 | 1st | |

*From stop codon

**Table S2.** RBS calculator predictions compared to observed measurements.

Candidate ribosome binding sequences (RBS), including their Shine Dalgarno (SD) sequences, were predicted using the RBS calculator [2] that both identifies and scores possible translation initiation sites, based on the 30S binding model for *de novo* translation initiation. The *de novo* initiation predictions showed no significant correlation with the observed GFP levels ($r^2$=0.08), with the levels of expression observed being generally more substantial than the predictions. This strengthens the argument that the expression of the distal operon gene encoding GFP to be mainly the result of re-initiation and not *de-novo* initiation.

| Clone | $\Delta G_{fold}$ | Best re-initiation start codon candidate(s) | Start codon position, relative to stop codon | Codon rank [1] | Observed translation rate [AU] | Predicted translation rate [AU] [2] | RBS binding energy $\Delta G_{total}$ [kcal/mol] [2] |
|---|---|---|---|---|---|---|---|
| **29** | -8.9 | I (AUU) | +8 | 6th | 32±4 | 0 | NA |
| **33** | -9.4 | V (GUG) | +8 | 2nd | 48±5 | 1.34 | 15.16 |
| **52** | -1.3 | M (AUG) | +5 | 1st | 344±12 | 90.75 | 5.80 |
| **56** | -8.5 | V (GUG) | +6 | 2nd | 62±4 | 6.31 | 11.72 |
| **57** | -6.5 | L (UUG) | +8 | 3rd | 43±7 | 0.49 | 17.40 |
| **62** | -6.7 | L (CUG) | +9 | 4th | 39±4 | 0 | NA |
| **71** | -4.9 | L (UUG) | +6 | 3rd | 98±4 | 13.79 | 9.98 |
| **91** | -2.5 | L (UUG) | +9 | 3rd | 103±4 | 0.96 | 15.91 |
| **96** | -2.1 | V (GUG) | +8 | 2nd | 197±7 | 5.76 | 11.92 |
| **101** | -6.0 | L (UUG) | +6 | 3rd | 74±24 | 35.90 | 7.86 |
| **110** | -8.2 | M (AUG) | +8 | 1st | 57±4 | 0.66 | 16.72 |
| **111** | -13.7 | I (AUU) | +5 | 6th | 43±3 | 0 | NA |
| **202** | -11.3 | V (GUG) | +4 | 2nd | 38±2 | 0.33 | 18.29 |
| **203** | -7.6 | L (UUG) | +7 | 3rd | 33±6 | 2.10 | 14.16 |
| **207** | -1.8 | M (AUG) | +3 | 1st | 526±53.5 | 16.52 | 9.58 |
| **208** | -1.9 | M (AUG) | +7 | 1st | 140±19 | 32.12 | 8.11 |
| **209** | -4.7 | M (AUG) | +7 | 1st | 274±8 | 479.88 | 2.10 |
| **212** | -5.9 | M (AUG) | +3 | 1st | 137±5 | 62.34 | 6.63 |
| **214** | -2.8 | M (AUG) | +3 | 1st | 478±17 | 11.74 | 10.34 |
| **216** | -0.8 | M (AUG) | +5 | 1st | 193±17 | 150.90 | 4.67 |
| **220** | -3.9 | M (AUG) | +6 | 1st | 201±16 | 1011.93 | 0.44 |
| **222** | -6.6 | M (AUG) | +3 | 1st | 211±13 | 1.78 | 14.53 |
| **225** | -7.5 | L (UUG) | +3 | 3rd | 78±4 | 1.61 | 14.76 |
| **226** | -9.2 | L (UUG) | +5 | 3rd | 56±6 | 0.70 | 16.59 |
| | | M (AUG) | +9 | 1st | | 3.31 | 13.16 |
| **230** | -5.0 | M (AUG) | +8 | 1st | 41±6 | 3.72 | 12.90 |
| **232** | -7.6 | M (AUG) | +6 | 1st | 36±5 | 122.95 | 5.12 |
| **233** | -8.4 | V (GUG) | +8 | 2nd | 25±5 | 0.13 | 20.39 |
| **235** | -9.5 | L (CUG) | +5 | 4th | 29±5 | 0 | NA |
| **236** | -7.9 | L (UUG) | +3 | 3rd | 65±4 | 0.28 | 18.63 |
| **238** | -8.4 | V (GUG) | +4 | 2nd | 86±5 | 1.54 | 14.86 |
| **244** | -3.5 | M (AUG) | +4 | 1st | 360±5 | 272.86 | 3.35 |
| **245** | -5.1 | M (AUG) | +7 | 1st | 222±11 | 710.51 | 1.23 |
| **249** | -1.9 | M (AUG) | +7 | 1st | 391±18 | 98.94 | 5.61 |

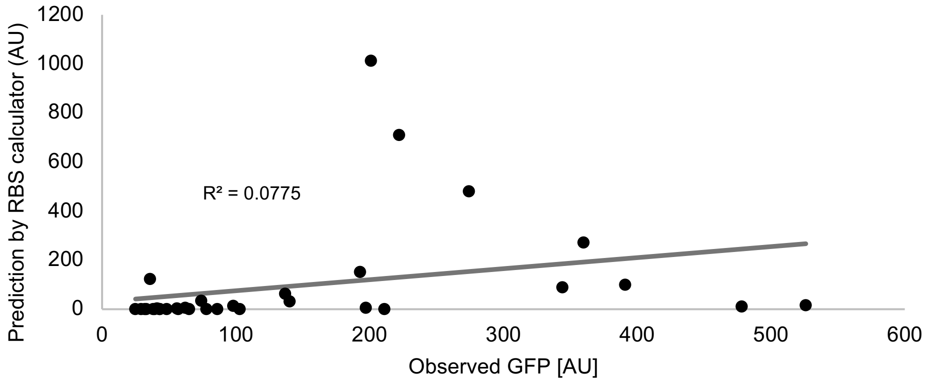


**Table S2-Fig. 1:** Correlation between observed GFP levels and those predicted upon *de novo* initiation using the RBS calculator.

**Table S3.** Statistical parameters for all tests.

| Test for result in figure | Test type | Alternative | Sampling position relative to stop codon[nt] | Name of sample one | N of sample two | Name of sample two* | N of sample two* | Central tendency [kCal/mol/ window] | Test statistic score | P-value | Notes |
|---|---|---|---|---|---|---|---|---|---|---|---|
| **Fig. 1d** | Mann-Whitney | Two-sided | N/A | P3 | 266,556 | P4 | 112,666 | P3= -6.440 P4= -6.307 (means) | | 0.0017 | |
| **Fig. 1d** | Mann-Whitney | One-sided: greater | N/A | P4 | 112,666 | P5 | 24,104 | P5= -5.996 (mean) | U= 1.4e9 | $<10^{-30}$ | |
| **Fig. 1d** | Mann-Whitney | One-sided: greater | N/A | P5 | 24,104 | P6 | 147,898 | P6= -5.895 (mean) | U= 1.8e9 | $2\times10^{-6}$ | |
| **Fig. 1d** | Mann-Whitney | One-sided: greater | N/A | P6 | 147,898 | P7 | 164,303 | P7=-5.809 (mean) | U= 1.2e10 | $3\times10^{-14}$ | |
| **Fig. 1d** | Mann-Whitney | One-sided: greater | N/A | P7 | 164,303 | P8 | 259,634 | P8= -5.712 (mean) | U= 2.2e10 | $<10^{-30}$ | |
| **Fig. 2b** | Wilcoxon | Two-sided | +5 | Sample | 82,360 (4,118 genes) | N/A | N/A | -2.37 (mean) | W=1.1e10 | $<10^{-30}$ | Was compared under the RTS model (see methods) |
| **Fig. 2d** | Mann-Whitney | Two-sided | +4 | High protein abundance | 24,440 | Low and medium protein abudance | 57,000 | -2.0 (median) | U=5.8e8 | $<10^{-30}$ | |
| **Fig. 3b** | Wilcoxon | Two-sided | 0 | <25 | 1,537 | N/A | N/A | 0.4 (mean) | W= 7.5e5 | $5\times10^{-19}$ | |
| **Fig. 3b** | Wilcoxon | Two-sided | 0 | ≥25 | 2,581 | N/A | N/A | -4.0 (mean) | W= 5.9e5 | $<10^{-30}$ | |
| **Fig. 3c** | Mann-Whitney | Two-sided | 0 | ≥25 | 51,620 | <25 | 30,920 | -2.0 (median) | U= 4.8e8 | $<10^{-30}$ | |
| **Fig. 4a** | Mann-Whitney | Two-sided | +3 | Last genes | 45,500 | Not last genes | 33,740 | -2.3 (median) | U= 5.0e8 | $<10^{-30}$ | |

*Two-sample testing

5

**Figure S2 – Quantitative PCR of synthetic operon mRNA levels in 5 clones:**

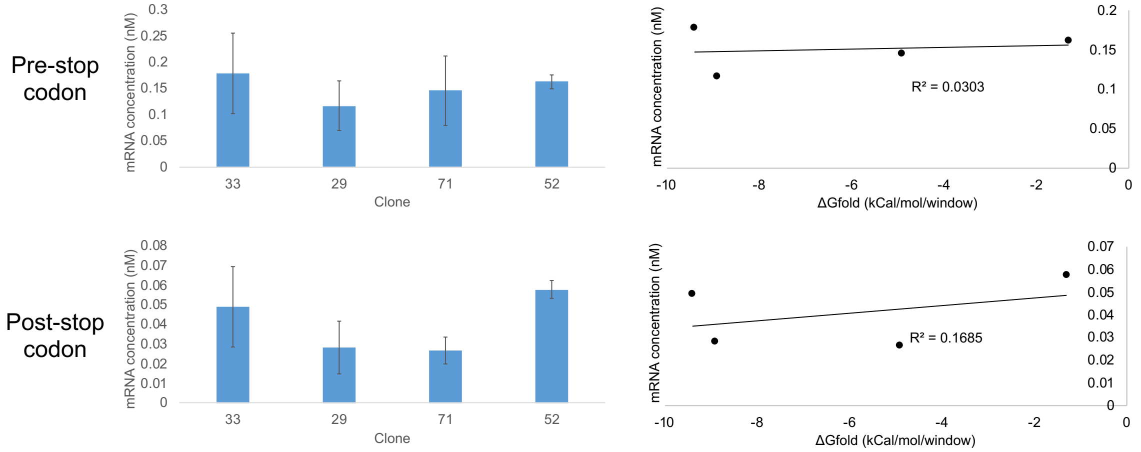


**Fig. S2.** mRNA abundance measured by 2 experimental repeats of qPCR, each with 3 replications of 4 select clones with different $\Delta G_{fold}$ (Bar graphs), error bars represent the STD of the mean. No significant correlation was noted between $\Delta G_{fold}$ of the variable region in several pRNXG clones and mRNA abundance in *E. coli* MG1655 (scatter plots). This was confirmed with amplicons of regions up-stream (pre-stop codon) and down-stream (post-stop codon) of the stop codon.

### Figure S3 – RFP expression from different synthetic operon clones:

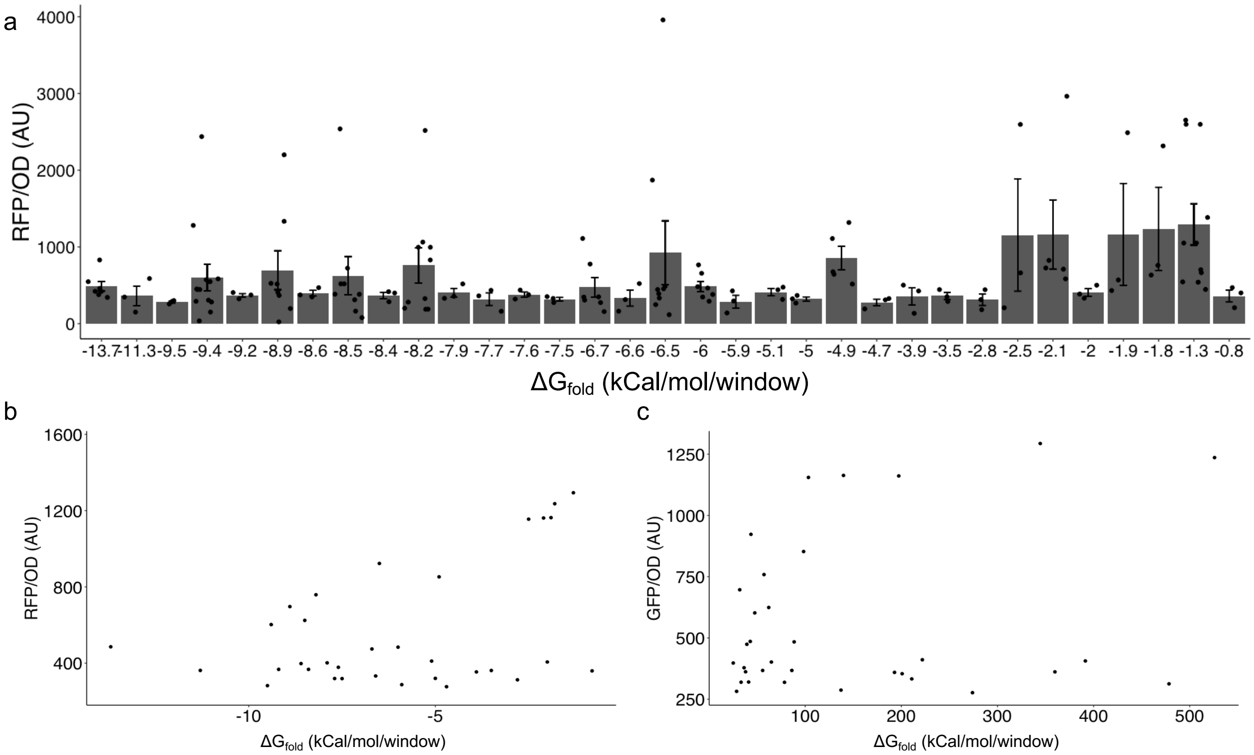


**Fig. S3. a**) Expression levels of RFP normalized to $OD_{600}$ measured by RFP fluorescence; error bars represent standard error of experimental repeats (n≥3 for all clones). **b**) Correlation between RFP fluorescence levels and $\Delta G_{fold}$. No significant correlation was observed (Spearman correlation=-0.19, S=7,118, n=33, p-val=0.29). **c**) Dependence between GFP and RFP expression levels of the synthetic operon. No significant correlation was observed (Spearman correlation=0.08, S=5,528 p-val=0.67).

**Figure S4 - ΔLFE of the three stop codons in *E. coli* and 128 other bacterial strains:**

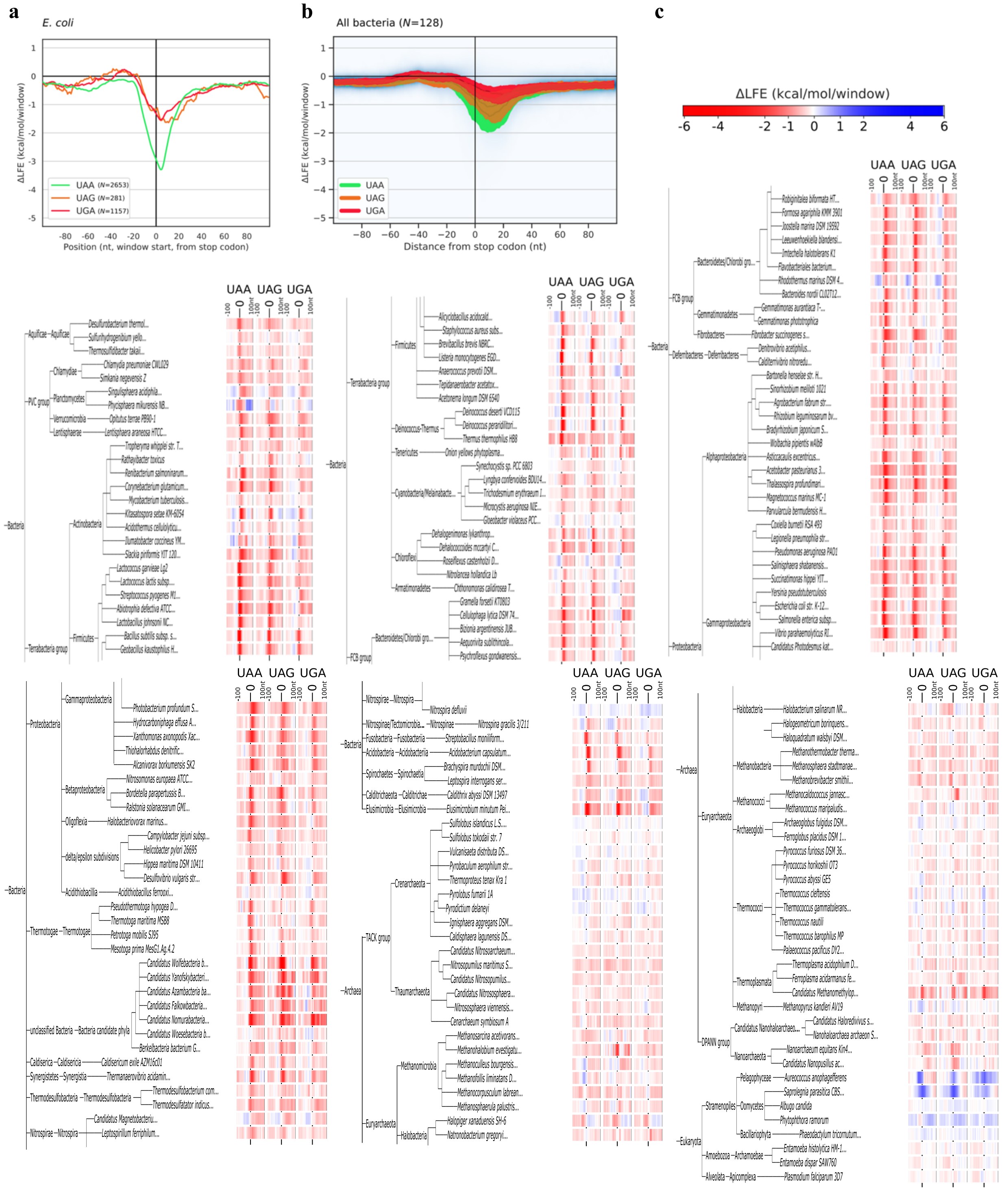


**Fig. S4 a**) All three *E. coli* stop codons were tested independently for RTS presence and strength. While the RTS was present after all stop codons, differences were observed. **b**) Where examined, RTS presence and stop codon-related differences in 128 bacterial species were, on average, consistent with what was observed in *E. coli*. In addition, ΔLFE heatmaps depicting the 100 nucleotide-long regions around stop codons across the tree of life for each of the three stop codons (warm colors: stronger folding than expected; cool colors: weaker folding than expected) was drawn. **c**) 128 bacterial, 49 archaeal, and 8 eukaryotic species were examined. The two latter domains presented mixed results.

**Figure S5 – RTS presence across of all kingdoms of life (all stop codons aggregated):**

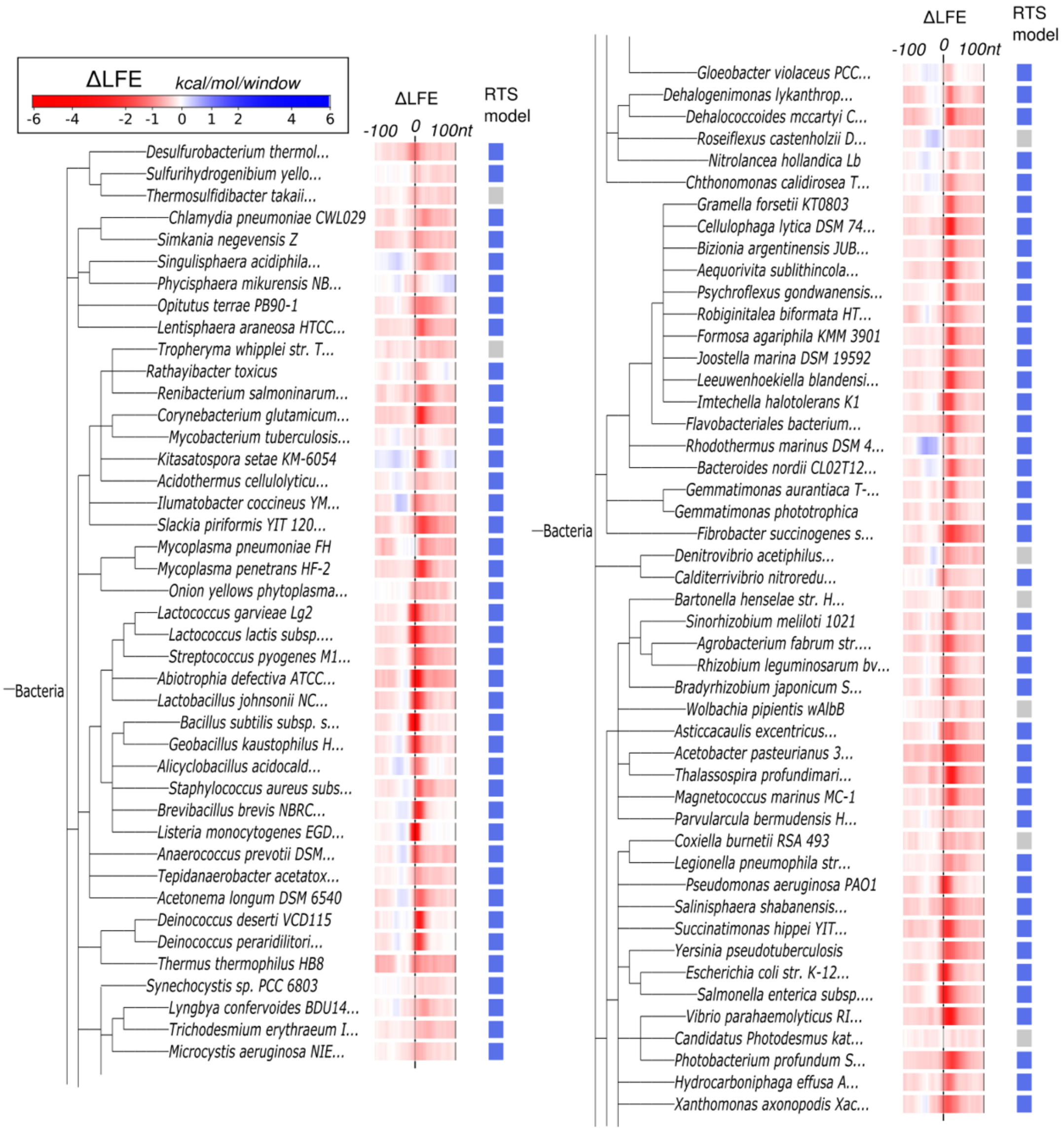

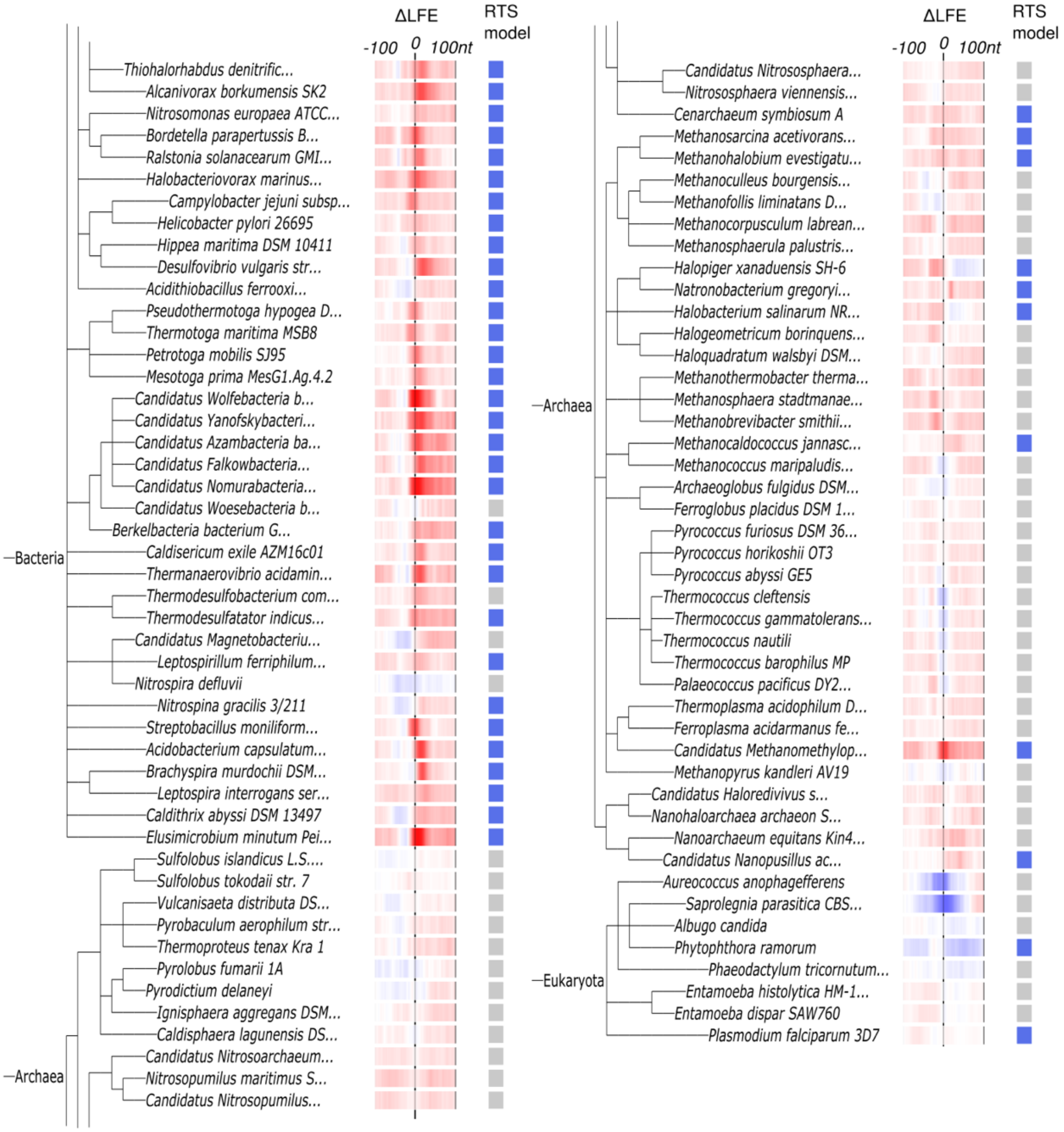


**Fig. S5.** The ΔLFE landscape was examined in 128 bacteria, 59 archaea, and 8 eukaryotes. The ΔLFE landscape was depicted as a heatmap of 100 nucleotide-long region around stop codons in species belonging to domains comprising the three branches of the tree of life (warm colors: stronger folding than expected; cool colors: weaker folding than expected). Using the RTS model (see Methods), we assessed the presence or absence of the RTS. The results revealed that 122/128 (95.3%) of bacteria, 12/49 (24.5%) of archaea and 2/8 (25.0%) of eukaryotes present an apparent RTS, although the sample sizes of the two latter groups are too small and the RTS signal is too weak and unreliable to draw any conclusions at this time.

**Figure S6 – Densitometric analysis of Western blots:**

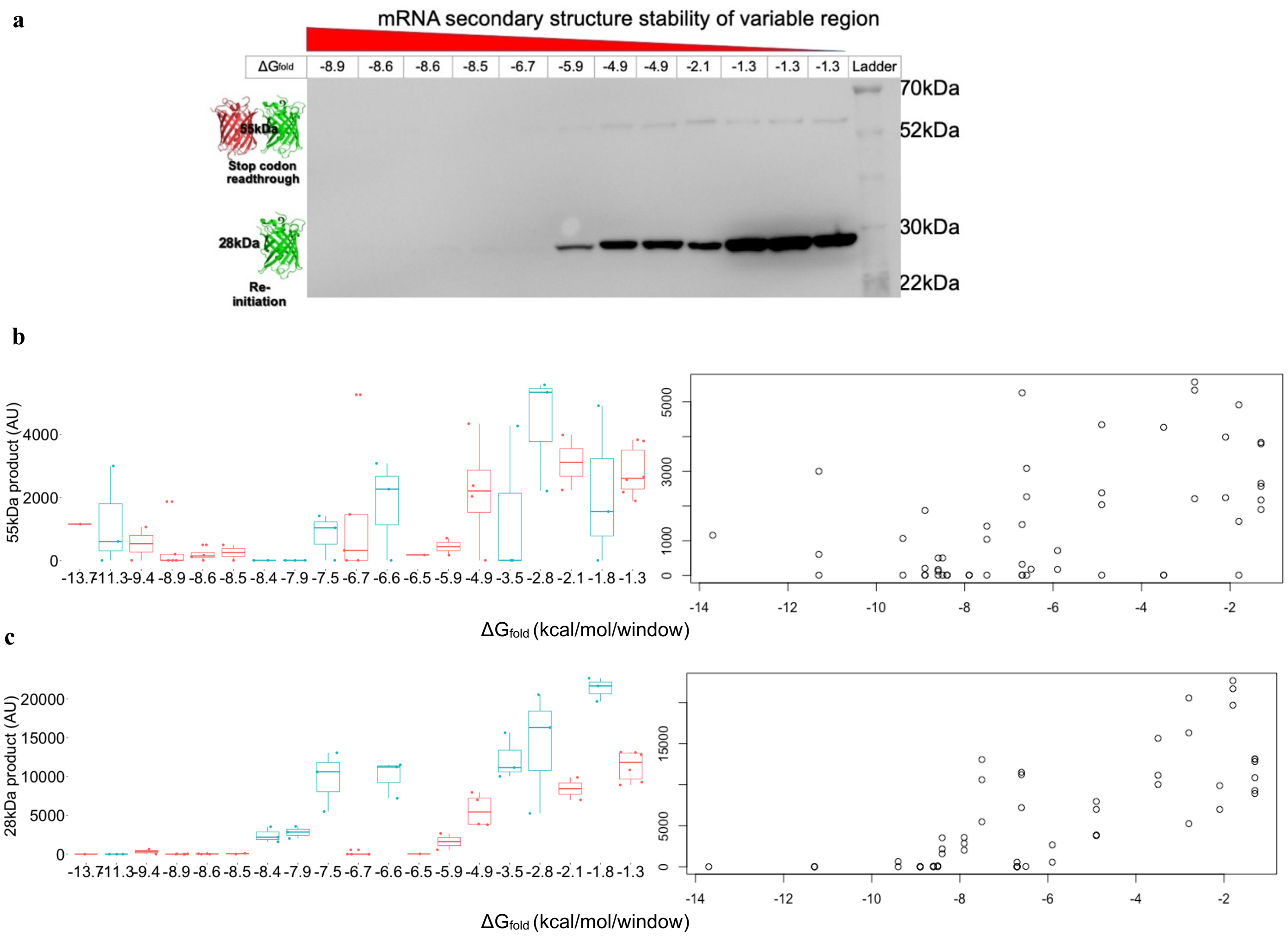


**Fig. S6. a**) Anti-His tag Western blot of random clones. For the randomly selected clones (red) and for the clones with an AUG start codon beginning at positions +3 or +4 (cyan), both **b)** the 55 kDa RFP-GFP product resulting from stop codon read-through, and **c)** the 28 kDa GFP product resulting from *de novo* initiation or re-initiation were measured using densitometry of the pRXNG clones in *E. coli* MG1655. The results were aggregated experimental repeats of each close as a box-plot for each clone (left) and as scatterplots for correlation analyses. In the scatterplots, each data point represents one experimental anti-His tag Western blot repeat of a clone with the indicated calculated $\Delta G_{fold}$. The 28 kDa GFP product accounts for 91% of the correlation between $\Delta G_{fold}$ and the total amount of GFP expressed by the different clones (omega squared test, $\omega^2$=0.91). Moreover, correlation with $\Delta G_{fold}$ was maintained for GFP (Spearman correlation, ρ=0.80, n=58, S=6479, p-val<$10^{-13}$) and also, albeit to a lesser degree, with the 55 kDa read-through product (Spearman correlation ρ=0.50, n=58, S=16326, p-val<$10^{-4}$).

### Figure S7 – Mass spectra of different clones:

Clone 52:

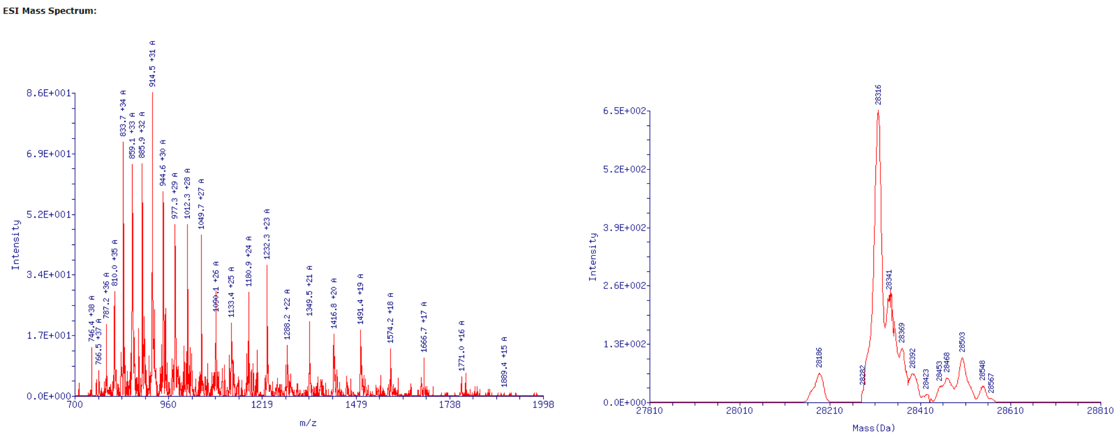


Clone 71:

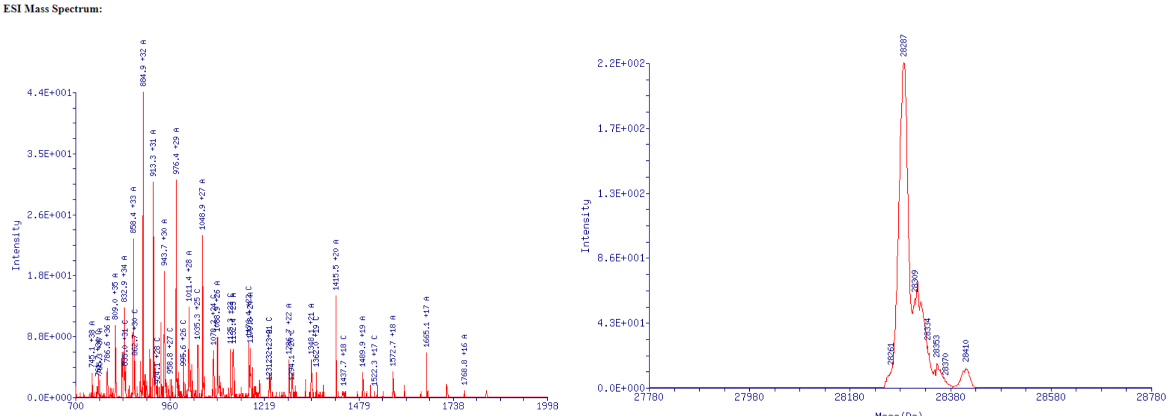


Clone 91:

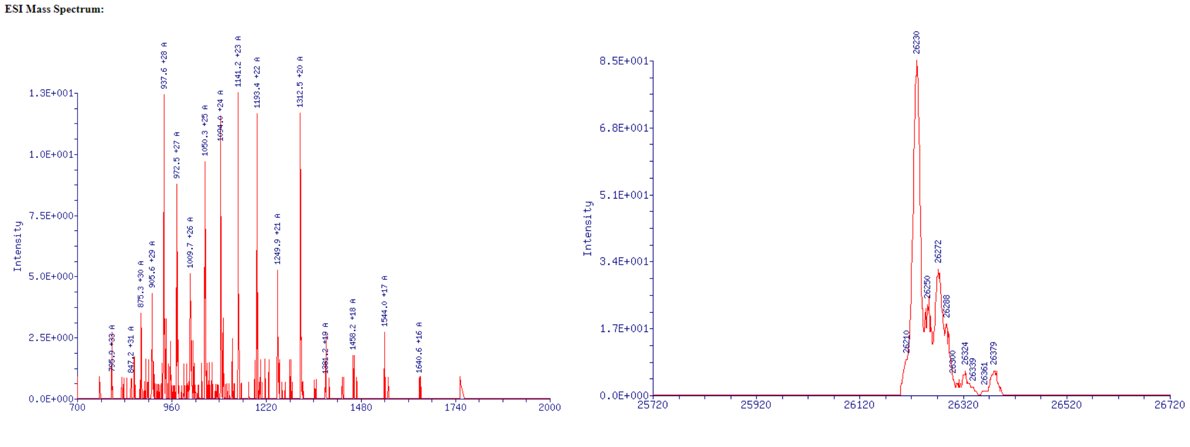

Clone 96:

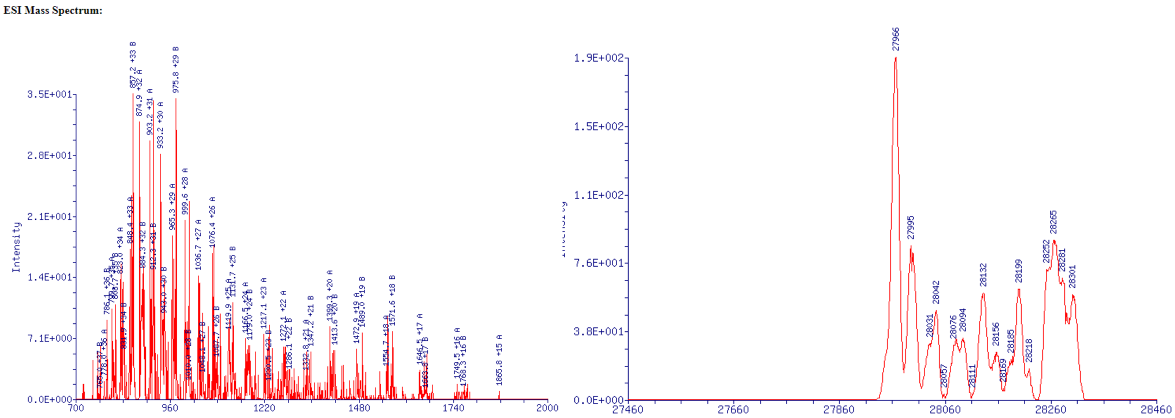


Clone 110:

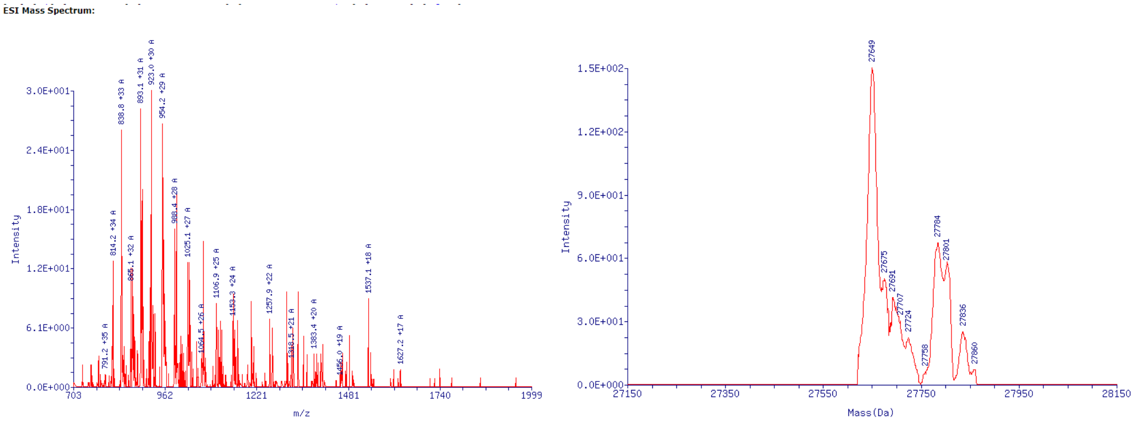


Full read-through product (entire dual reporter construct):

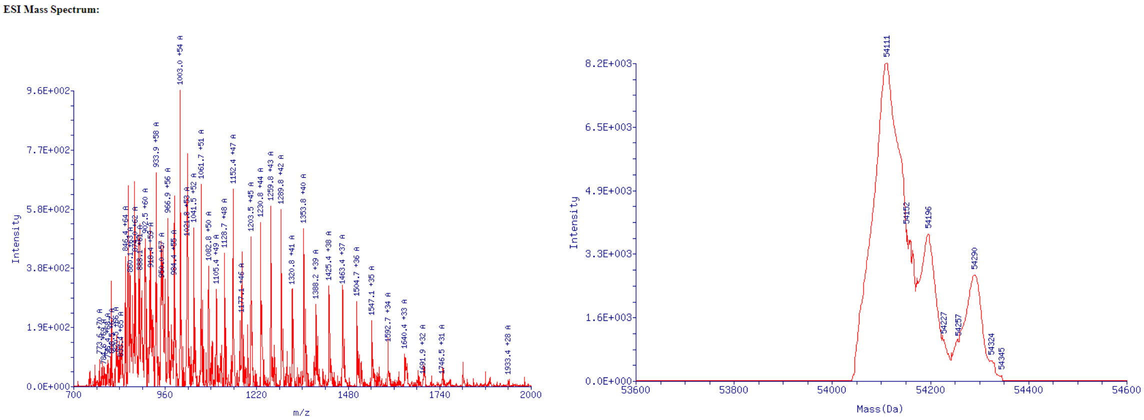


**Fig. S7.** Five clones expressing sufficient levels of the ~28 kDa GFP product and a representative read-through product (with the UAG stop codon mutated to encode tyrosine) were purified using nickel affinity columns and subjected to mass spectrometry to identify the start codon. These involved comparisons of calculated masses generated by the clone-specific sequence and the measured mass of the protein. Left panels depict the raw MS results, while the right panels depict de-convoluted data obtained using Promass software. In the manuscript, we report the primary product of each clone. However, we cannot exclude or accurately assess the possibility of multiple possible initiation sites with different efficiencies.

### Figure S8 – Correlation between $\Delta G_{fold}$ and GFP levels without and with Release Factor 1 (RF1):

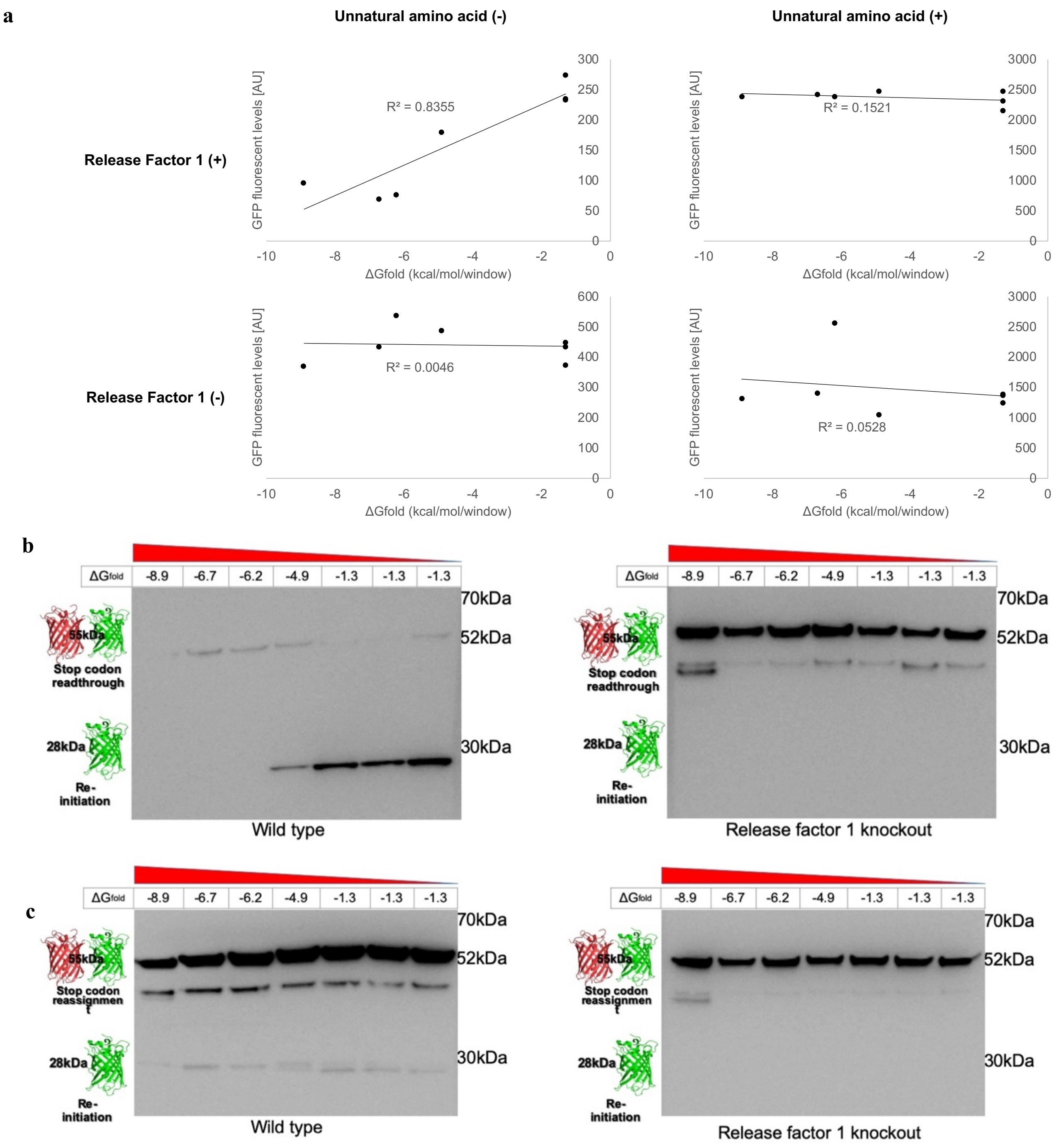


**Fig. S8. a**) Comparison of GFP expression, measured by fluorescence, between *E. coli* C321.ΔprfA EXP and MG1655, both transformed with the pEVOL pylRS genetic code expansion system and 5 PRXNG clones with different $\Delta G_{fold}$. Each data point represents the average of n=3 experimental replicates. **b**) Uncropped anti-His-tag Western blots presented in Fig. 3G. of the same 5 clones mentioned above, **c**) Uncropped gels presented in Fig. 3H. The bands below the RFP-GFP product (with a size of ~50kDa) are the his-tagged pyrrolysyl synthetase (*pylRS)* gene from the co-transformed pEVOL plasmid which is used for genetic code expansion is transformed.

**Figure S9 - Analysis of operonic position effect on RTS presence with/without a down-stream AUG start codon:**

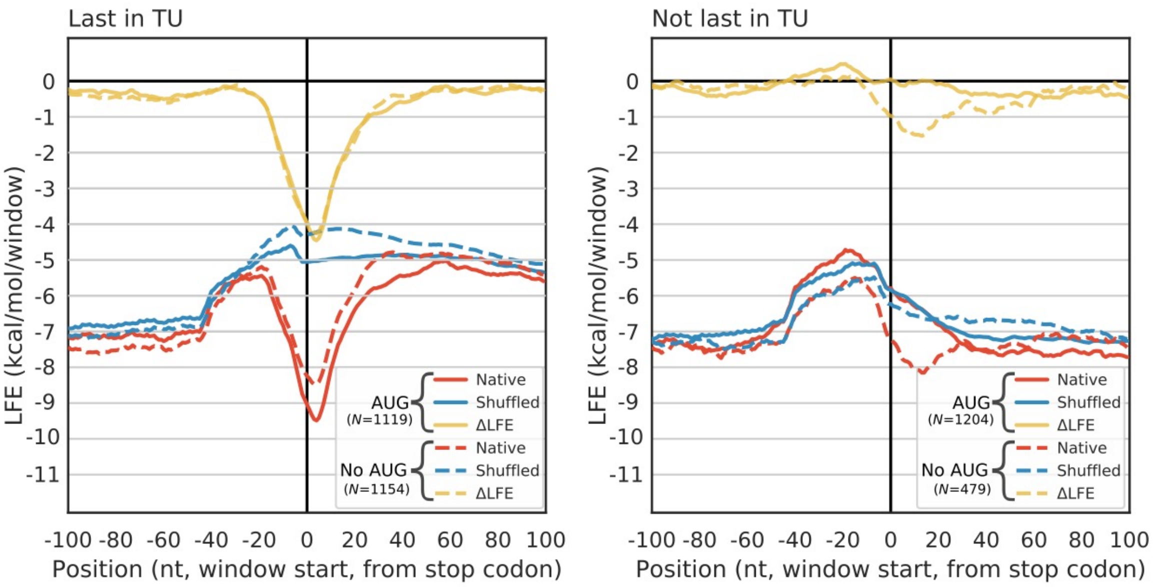


**Fig. S9.** Terminal operonic genes either with or without an AUG start codon in-frame of the down-stream CDS in the 50 nt downstream of a stop codon. Right panel: Mid-operonic genes either with or without an AUG start codon in-frame of the down-stream CDS in the 50 nt downstream of a stop codon. We examined differences between two groups of genes, namely those assuming the last position in an operon (i.e., terminal genes) **(left)** versus all other operon genes (i.e., non-terminal genes) **(right)**. Each group was further divided according to the presence of an in-frame AUG start codon within 50 nt downstream of the stop codon or the absence of a start codon. Such divisions revealed that in terminal genes, where translation insulation is expected in all cases, significant selection for an RTS was observed, regardless of the presence or absence of a down-stream start codon. Conversely, in mid-operon genes, selection for RTSs in the group with the start codon, where re-initiation is expected, is not higher than random. In the second group, where re-initiation is not desired as no in-frame AUG start codon exists, significant selection for RTSs was observed.

**Figure S10 - Controlling for an RTS link to transcription termination:**

We controlled for the fact that a stable mRNA structure down-stream of a stop codon could be functionally related to transcription termination since rho-independent transcription terminators can form stable mRNA hairpins. Therefore, to distinguish the role of the RTS in regulating translation re-initiation from transcription termination, all 871 known or suspected genes that terminate with a rho-independent terminator sequence[3] were removed from the analysis (Fig. S10, left). The RTS signal remained (Wilcoxon test, p-val<$10^{-16}$). The reduction in the effect is probably due to the fact that Rho-independent terminators affected the analysis by biasing the sequences ~40-60 nt downstream of the stop codon to more stable structures, thus interfering with our analysis around the stop codon, as the window size used was 40 nt. To further demonstrate the absence of a link between the RTS and transcription termination, two subsets of terminal and monocistronic genes were analyzed according to their experimentally measured 3' UTR lengths[4] (Fig. S10, right), with one group presenting short 3' UTRs (<50 nt) and the other possessing long 3' UTRs (>50 nt). Were the RTS signal linked to transcription termination, one would expect to see the RTS signal closer to the stop codons in the former and further away from the stop codon in the latter. However, no change in the position or magnitude of the RTS was observed. These analyses, taken together, demonstrate that the RTS is not linked to transcription termination.

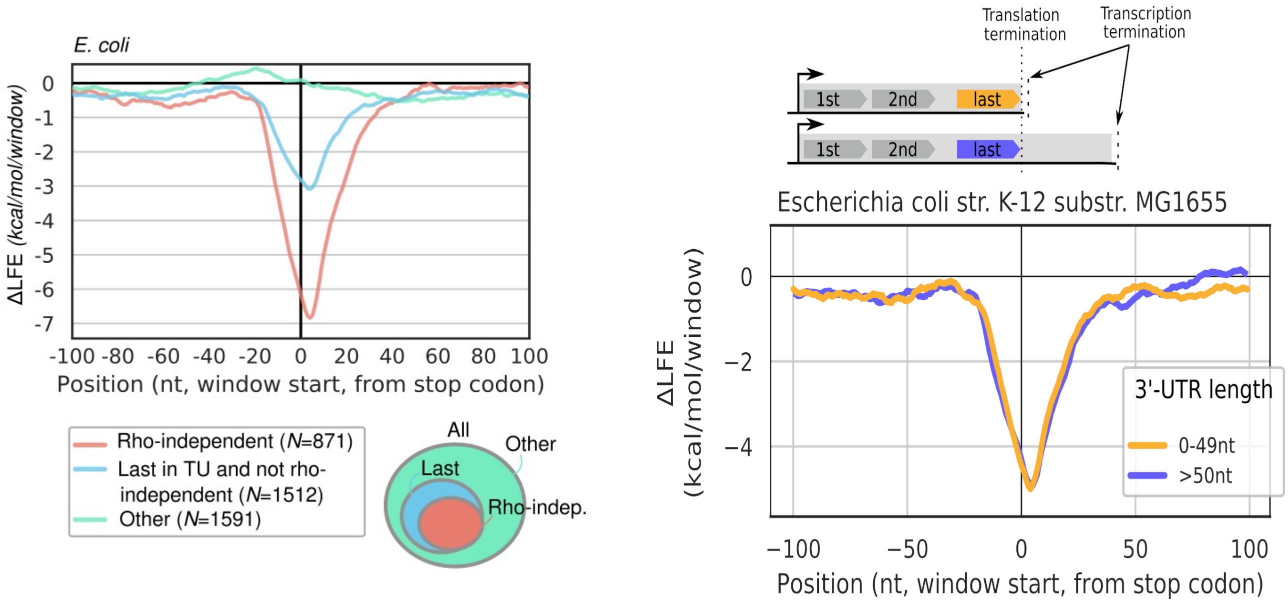


**Fig. S10. Left panel)** Analysis of *E. coli* genes grouped by transcription termination mechanism shows that folding bias cannot be explained by the presence of rho-independent terminators. Red, genes with rho-independent terminators. Blue, genes that are last in their transcription units (TU) but do not have rho-independent terminators. Green, all other genes. Lines represent ΔLFE, computed as described in the Methods section. Annotation of rho-independent genes based on WebGesTer-DB. Annotation of TU positions based on the ODB4 database. **Right panel)** The RTS signal shows no change between groups of genes with short (<50 nt) or long (>50 nt) 3' UTRs.

**Figure S11- Probability of having a start codons downstream of a stop codon without selection:** When considering the evolution of translation re-initiation, two solutions to avoid un-intended re-initiations when this is deleterious (for example, after the last gene of a poly-cistronic mRNA) are possible. The first involves depleting all efficient start codons. However, this is not optimal for three reasons: i) Even inefficient start codons could lead to basal expression by re-initiation; ii) ribosomes would wastefully spend time scanning for start codons which are depleted, resulting in a fitness cost; and iii) the probability of efficient start codons (one of the 6 most efficient[1]) on a random 3'UTR sequence is >0.9 (Fig. S11A) if considering the median *E. coli* 3' UTR length of 50 nucleotides (Fig. S11D). Moreover, the selection on the 3' UTR would have to be extremely high to counter the ~17% chance of an efficient start codon appearing after each single nucleotide mutation (Fig. S11B). This constraint is further compounded by consecutive mutations (Fig. S11C). To assess the length of *E. coli* 3'UTRs , we utilized RNA-seq data [4]. The data revealed that in *E. coli,* the average 3' UTR length is 76 nucleotides, with the median length being 50 nucleotides, a sufficient length to harbor significant mRNA secondary structure, and require stringent selection to avoid start codon-generated mutations.

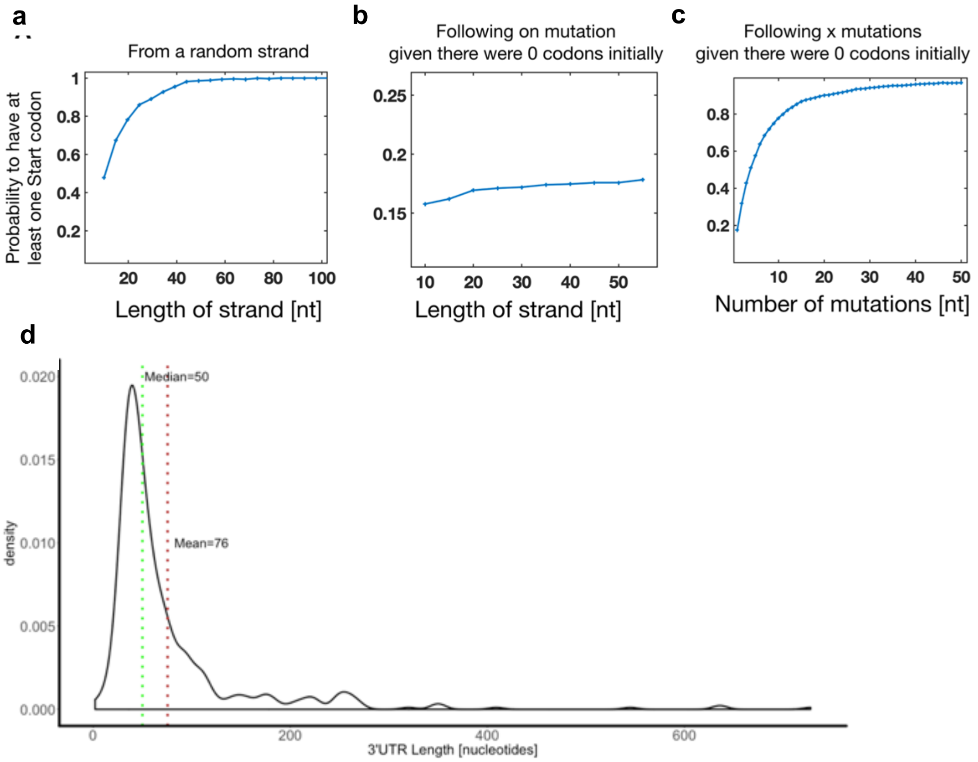


**Fig. S11. a**) The probability of having at least one efficient start codon (ATG, GTG, TTG, CTG, ATA, ATT) by chance as a function of DNA length. **b**) The probability that a sequence with no efficient start codon will generate an efficient start codon after a one nucleotide mutation as a function of strand length (Juke and Cantor, one parameter mutation model). **c**) The probability of having at least one efficient start codon through consecutive mutations on a fixed, 50 base pair-long DNA stretch. **d**) Density plot of mapped *E. coli* 3' UTR lengths in the RegulonDB database[4] (470 transcriptions units).

**Supplementary References**


1. Hecht, A. *et al.* Measurements of translation initiation from all 64 codons in E. coli. *Nucleic Acids Res.* **45**, 3615–3626 (2017).
2. Espah Borujeni, A. & Salis, H. M. Translation Initiation is Controlled by RNA Folding Kinetics via a Ribosome Drafting Mechanism. *J. Am. Chem. Soc.* **138**, 7016–7023 (2016).
3. Mitra, A., Kesarwani, A. K., Pal, D. & Nagaraja, V. WebGeSTer DB-A transcription terminator database. *Nucleic Acids Res.* **39**, 129–135 (2011).
4. Gama-Castro, S. *et al.* RegulonDB version 9.0: High-level integration of gene regulation, coexpression, motif clustering and beyond. *Nucleic Acids Res.* **44**, D133–D143 (2016).
